# SARS-CoV-2 3CLpro whole human proteome cleavage prediction and enrichment/depletion analysis

**DOI:** 10.1101/2020.08.24.265645

**Authors:** Lucas Prescott

## Abstract

A novel coronavirus (SARS-CoV-2) has devastated the globe as a pandemic that has killed more than 1,600,000 people. Widespread vaccination is still uncertain, so many scientific efforts have been directed toward discovering antiviral treatments. Many drugs are being investigated to inhibit the coronavirus main protease, 3CLpro, from cleaving its viral polyprotein, but few publications have addressed this protease’s interactions with the host proteome or their probable contribution to virulence. Too few host protein cleavages have been experimentally verified to fully understand 3CLpro’s global effects on relevant cellular pathways and tissues. Here, I set out to determine this protease’s targets and corresponding potential drug targets. Using a neural network trained on cleavages from 388 coronavirus proteomes with a Matthews correlation coefficient of 0.983, I predict that a large proportion of the human proteome is vulnerable to 3CLpro, with 4,460 out of approximately 20,000 human proteins containing at least one putative cleavage site. These cleavages are nonrandomly distributed and are enriched in the epithelium along the respiratory tract, brain, testis, plasma, and immune tissues and depleted in olfactory and gustatory receptors despite the prevalence of anosmia and ageusia in COVID-19 patients. Affected cellular pathways include cytoskeleton/motor/cell adhesion proteins, nuclear condensation and other epigenetics, host transcription and RNAi, ribosomal stoichiometry and nascent-chain detection and degradation, coagulation, pattern recognition receptors, growth factors, lipoproteins, redox, ubiquitination, and apoptosis. This whole proteome cleavage prediction demonstrates the importance of 3CLpro in expected and nontrivial pathways affecting virulence, lead me to propose more than a dozen potential therapeutic targets against coronaviruses, and should therefore be applied to all viral proteases and subsequently experimentally verified.

## Introduction

Coronaviruses are enveloped, positive-sense, single-stranded RNA viruses with giant genomes (26-32 kb) that cause diseases in many mammals and birds. Since 2002, three human coronavirus outbreaks have occurred: severe acute respiratory syndrome (SARS) in 2002-2004, Middle East respiratory syndrome (MERS) from 2012 to present, and coronavirus disease 2019 (COVID-19) from 2019 to present. The virus that causes the latter disease, SARS-CoV-2, was first thought to directly infect the lower respiratory epithelium and cause pneumonia in susceptible individuals. The most common symptoms include fever, fatigue, nonproductive or productive cough, myalgia, anosmia, ageusia, and shortness of breath. More recently, however, correlations between atypical symptoms (chills, arthralgia, diarrhea, conjunctivitis, headache, dizziness, nausea, severe confusion, stroke, and seizure) and severity of subsequent respiratory symptoms and mortality have motivated researchers to investigate additional tissues that may be infected. One way to explain these symptoms and associated cellular pathways is to review enrichment and depletion in virus-host interaction networks, particularly those including the coronavirus proteases.

Angiotensin-converting enzyme 2 (ACE2), the main receptor for SARS-CoV-1 and −2, has been shown to be less expressed in lung than in many other tissues. Respiratory coronaviruses likely first infect the nasal epithelium and tongue[1] and then work their way down to the lung and/or up through the cribriform plate to the olfactory bulb, through the rhinencephalon, and finally to the brainstem.[2–5] Additionally, based on ACE2 expression and *in vitro* and *in vivo* models, multiple parts of the gastrointestinal tract (mainly small and large intestine, duodenum, rectum, and esophagus; less appendix and stomach) and accessory organs (mainly gallbladder, pancreas, liver[6, 7], salivary gland[8]; less tongue and spleen)[9], kidney,[10] male and female reproductive tissues,[11, 12] heart,[13] immune cells,[14, 15] and adipose tissue[16–18] may be infectible with corresponding symptoms and comorbidities.

Coronaviruses have two main open reading frames, orf1a and orf1b, separated by a ribosomal frameshift and resulting in two large polyproteins, pp1a and pp1ab, containing proteins including two cysteine proteases,[19] an RNA-dependent RNA polymerase, and other nonstructural proteins (nsp1-16). The main function of these proteases is to cleave the polyproteins into their individual proteins to form the transcription/replication complex, making them excellent targets for antiviral drug development.[20–23] The papain-like protease (PLpro) and 3 chymotrypsin-like protease (3CLpro) only have 3 and 11 cleavage sites, respectively, in the polyproteins, but it is reasonable to assume that both proteases may cleave host cell proteins to modulate the innate immune response and enhance virulence as in picornaviruses and retroviruses, such as human immunodeficiency virus (HIV).

PLpro is a highly conserved protein domain that has been shown to determine virulence of coronaviruses[24] and possess deubiquinating and deISGylating activity including cleaving interferon-stimulated gene 15 (ISG15) induced by interferon via the Janus kinases and signal transducer and activator of transcription proteins (JAK-STAT) pathway from ubiquitin-conjugating enzymes and potentially from downstream effectors.[25–29] PLpro deubiquination also prevents activating phosphorylation of interferon regulatory factor 3 (IRF3) and subsequent type-I interferon production,[30, 31] however the ubiquitinated leucine in human IRF3 is replaced by a serine in bats likely including *Rhinolophus affinus* (intermediate horseshoe bat), the probable species of origin of SARS-CoV-2.[32, 33]

3CLpro is also highly conserved among coronaviruses; SARS-CoV-2 3CLpro is 96.08% and 50.65% identical, respectively, to the SARS- and MERS-CoV homologs, the former with only 12 out of 306 amino acids substituted with all 12 outside the catalytic dyad or surrounding pockets.[34–36] Even the most distant porcine deltacoronavirus HKU15 3CLpro shares only 34.97% identity yet is similarly conserved in the these important residues. This conservation indicates that all these proteases are capable of cleaving similar sequences no matter the protease genus of origin. In addition to the 11 sites in the polyproteins, these proteases are known to cleave host proteins including STAT2[37], NF-kappa-B essential modulator (NEMO)[38], the nucleotide-binding oligomerization domain (NOD)-like receptor NLRP12, and TGF-beta activated kinase 1 (TAB1)[39] to modulate interferon signaling. Similar proteases have been studied in the other members of *Nidovirales*[40] and the related *Picornavirales*[41–45], with foot-and-mouth disease virus (FMDV) 3Cpro cleaving histone H3,[46, 47] poliovirus 3Cpro cleaving TFIID and TFIIIC,[48–52] and polio- and rhinovirus but not cardiovirus 3Cpro cleaving microtubule-associated protein 4 (MAP4).[53, 54] These results, however, have not been reproduced for SARS-CoV-2 yet, and STAT2, NEMO, NLRP12, TAB1, H3, TFIIIC, TFIID, and MAP4 are only a few of many cleaved proteins.

The high number of 3CLpro cleavages in coronavirus polyproteins has, however, allowed for sequence logos and resulting sequence rules and training of decision trees and neural networks (NN) for additional cleavage site prediction.[55–60] Notably, Kiemer et al.’s NN[59] based on Blom et al.’s equivalent picornaviral NN[60] was trained on 7 arbitrary coronavirus genomes, totaling 77 cleavages, and had a Matthews correlation coefficient (MCC) of 0.84, much higher than the traditional consensus pattern’s 0.37 for the same training set size. They predicted cleavage sites in select host proteins, namely the transcription factors CREB-RP, OCT-1, and multiple subunits of TFIID, the innate immune modulators interferon alpha-induced protein 6 (IFI6) and IL-1 receptor-associated kinase 1 (IRAK-1), the epithelial ion channels cystic fibrosis transmembrane conductance regulator (CFTR) and amiloride-sensitive sodium channel subunit delta (SCNN1D), the tumor suppressors p53-binding proteins 1 and 2 (although not p53 itself), RNA polymerase I and III subunits (RPA1 and RPC1), eukaryotic translation initiation factor 4 gamma 1 (eIF4G1), the cytoskeletal proteins MAP4 and microtubule-associated protein RP/EB members 1 and 3 (MAPRE1/3), and many members of the ubiquitin pathway (ubiquitin hydrolases USP1/4/5/9X/9Y/13/26 and suppressor of cytokine signaling 6 (SOCS6)).

Additionally, Yang’s decision trees[58] were trained on 4 amino acid sliding windows and substitution matrix similarity score-based embeddings, achieved MCCs up to 0.95, but were limited to only 18 coronavirus polyproteins. The embedding-derived non-orthogonality somewhat stabilized the prediction to small changes in sequence assuming the substitution matrix reflects how the cleavages evolve. Decision trees have the benefit of being symbolic and explainable but often predict suboptimally when presented with interpolated or extrapolated inputs, making alternative machine learning techniques more attractive for predicting human protein cleavage prediction. For example, Narayanan et al.[61] and later Singh et al.[62] demonstrated that neural networks outperform decision trees for HIV and hepatitis C virus (HCV) protease cleavage prediction. Additional mixed methods such as Li et al.’s nonlinear dimensionality reduction and subsequent support vector machine (SVM) are able to retain some of the benefits of both linear and nonlinear classifiers.[63] Rognvaldsson et al.[64, 65] argue that nonlinear models including neural networks should not be used for cleavage prediction, however the HIV dataset from Cai et al.[66] that they used and their expanded dataset only included 299 and 746 samples, respectively. Additionally, physiochemical or structural encodings have outperformed one-hot encoding (also called orthogonal encoding) for their small HIV datasets[67] and have moreover eliminated differences between linear and nonlinear classifiers in an equivalent HCV dataset with 891 samples.[68] To my knowledge no one has expanded the 3CLpro cleavage dataset to the point where nonlinearity becomes significant, investigated the entire human proteome for 3CLpro cleavages sites with any method, or performed enrichment analysis and classification of these affected proteins.

## Methods

### Data Set Preparation

A complete, manually reviewed human proteome containing 20,350 sequences (not including alternative isoforms) was retrieved from UniProt/Swiss-Prot (proteome:up000005640 AND reviewed:yes).[69]

Additional coronavirus polyprotein cleavages were collected from GenBank.[70] Searching for “orf1ab,” “pp1ab,” and “1ab” within the family *Coronaviridae* returned 388 different, complete polyproteins with 762 different cleavages manually discovered using the Clustal Omega multiple sequence alignment server.[71–73] All 4,268 balanced positive cleavages were used for subsequent classifier training in addition to all other uncleaved coronavirus sequence windows centered at glutamines (17,493) and histidines (11,421), totaling 33,182 samples.

### Cleavage Prediction

The NetCorona 1.0 server as in Kiemer et al.’s work[59], my reproductions of their sequence logo-derived rules and NN, and my improved sequence logo-based logistic regression and naïve Bayes classification and NNs were used for prediction of cleavage sites.[74] Some predicted cleavage sites were close enough to the N- and C-termini that the nine amino acid window input into the neural network was not filled. These sites with center glutamine residue less than four amino acids from the N-terminus or less than five amino acids from the C-terminus were omitted because although they may be within important localization sequences, their cleavage kinetics are likely significantly retarded by truncation.

### Enrichment Analysis

Protein annotation, classification, and enrichment analysis was performed using the Database for Annotation, Visualization, and Integrated Discovery (DAVID) 6.8.[75, 76] My training data, prediction methods, and results can be found on GitHub (https://github.com/Luke8472NN/NetProtease).

## Results

Here I assumed that SARS-CoV-2 3CLpro is capable of cleaving all aligned cleavages between the four genera of coronaviruses (*Alpha-*, *Beta-*, *Gamma-*, and *Delta-*) because variation in cleavage sequences is greater within polyproteins than between them (Figure 1) no matter the existence of protease/cleavage cophylogeny (Figure 2).[77]

**Figure 1:**
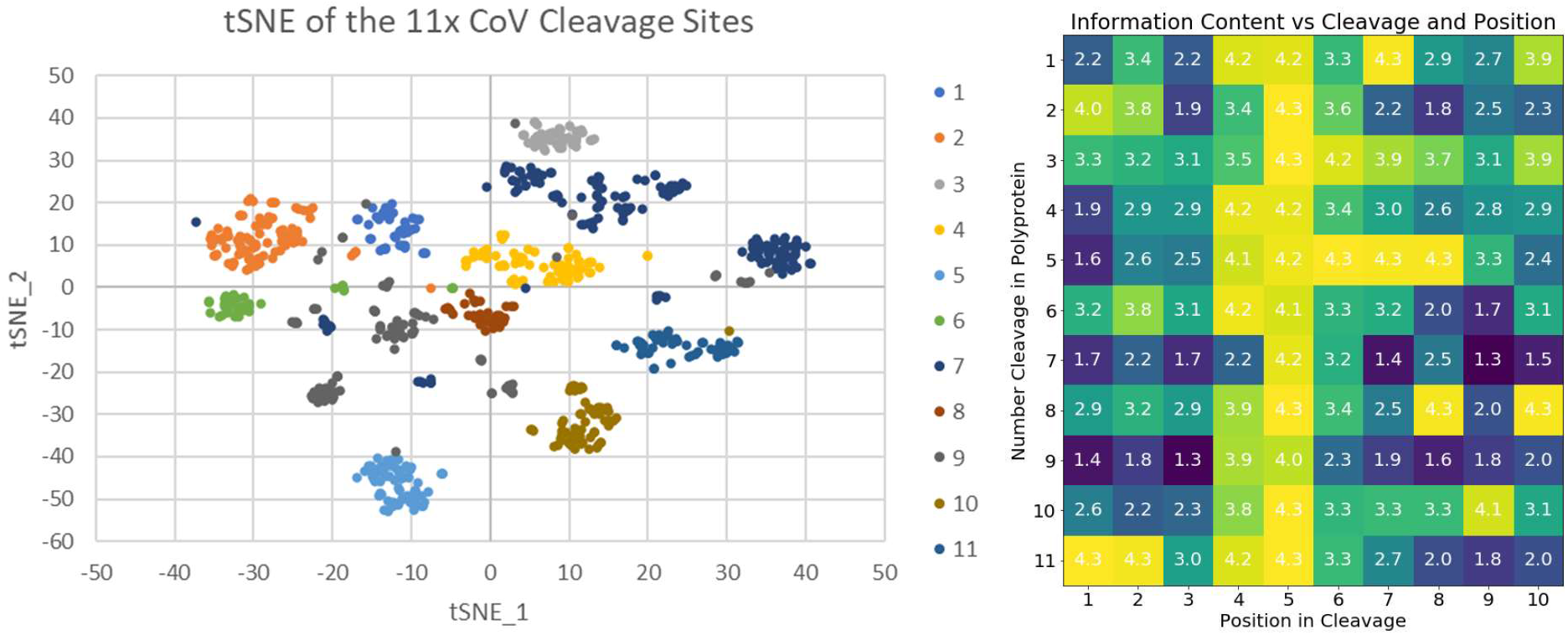
One-hot encoded t-distributed stochastic neighbor embedding (t-SNE)[78] and information content both demonstrate that cleavage variation within genomes is more important than variation between genomes.

**Figure 2:**
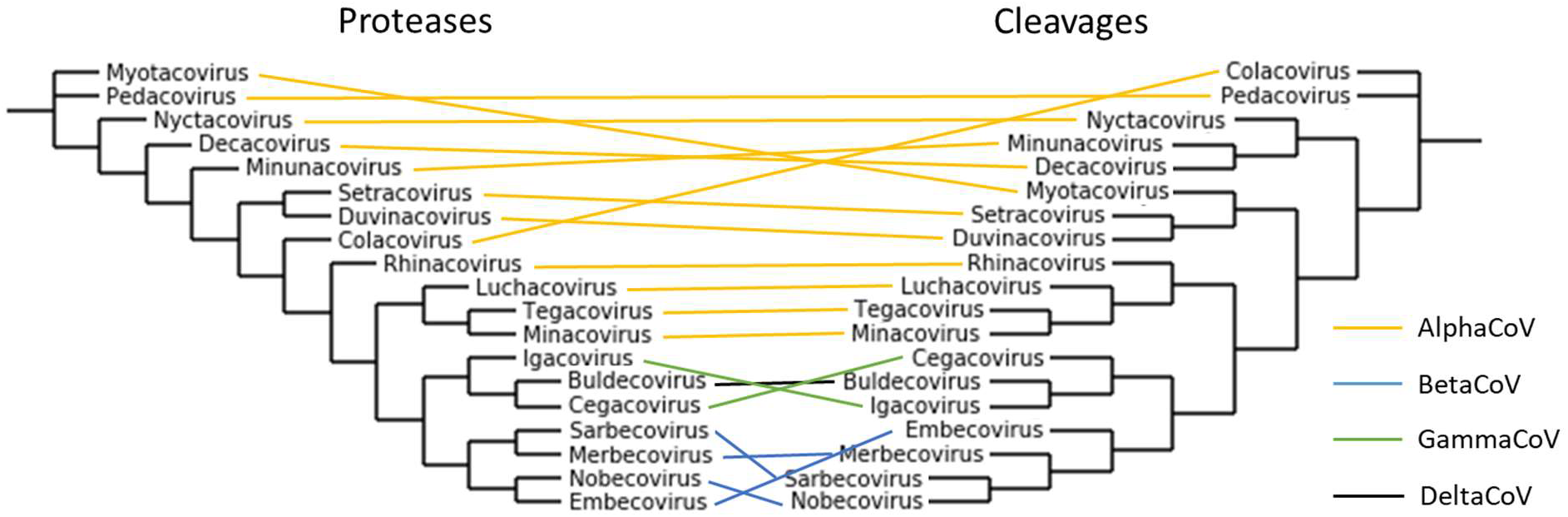
Unscaled subgenera-averaged tanglegram of 3CLpro and respective cleavages based on BLOSUM62 substitution matrix similarity scores with and without default affine gap penalties (opening 10 and extension 0.2).

Kiemer et al.’s seven genome sequence logo and multilayer perceptron structure with each amino acid one-hot encoded as a binary vector of length 20 (an input of 200 bits i.e. linearized 10 amino acids surrounding the cleavage) were both reproduced.[59] First, logistic regression was performed on the logit of the probability output of the sequence logo (as opposed to Chou et al.’s manual probability cutoff setting by maximizing an unbalanced measure of accuracy[79]) with a nonzero but optimally extremely small pseudocount and returned an MCC of 0.825 with 74.0% recall. Updating the sequence logo with all known cleavages improved its MCC to 0.936 with 94.8% recall (Figure 3). A naïve Bayes classifier was additionally constructed from both the positive and negative sequence logos and slightly improved the MCC to 0.947 with 95.7% recall. Figure 4 demonstrates correlations (represented as the mutual information variant known as total entropy correlation coefficients or symmetric uncertainties) between positions that are not captured by simple sequence logos and classifiers assuming independence.[80, 81] NNs, however, allow inclusion of 2D and higher-order correlations not easily visualizable and therefore often improve accuracy. Finally, in addition to information content, Figure 5 shows a charge-polarity-hydrophobicity scale with a lack of obvious trend or conservation reaffirming that one-hot encoding performs better than any physiochemical, lower-dimensional inputs when the training set is large enough.

**Figure 3:**
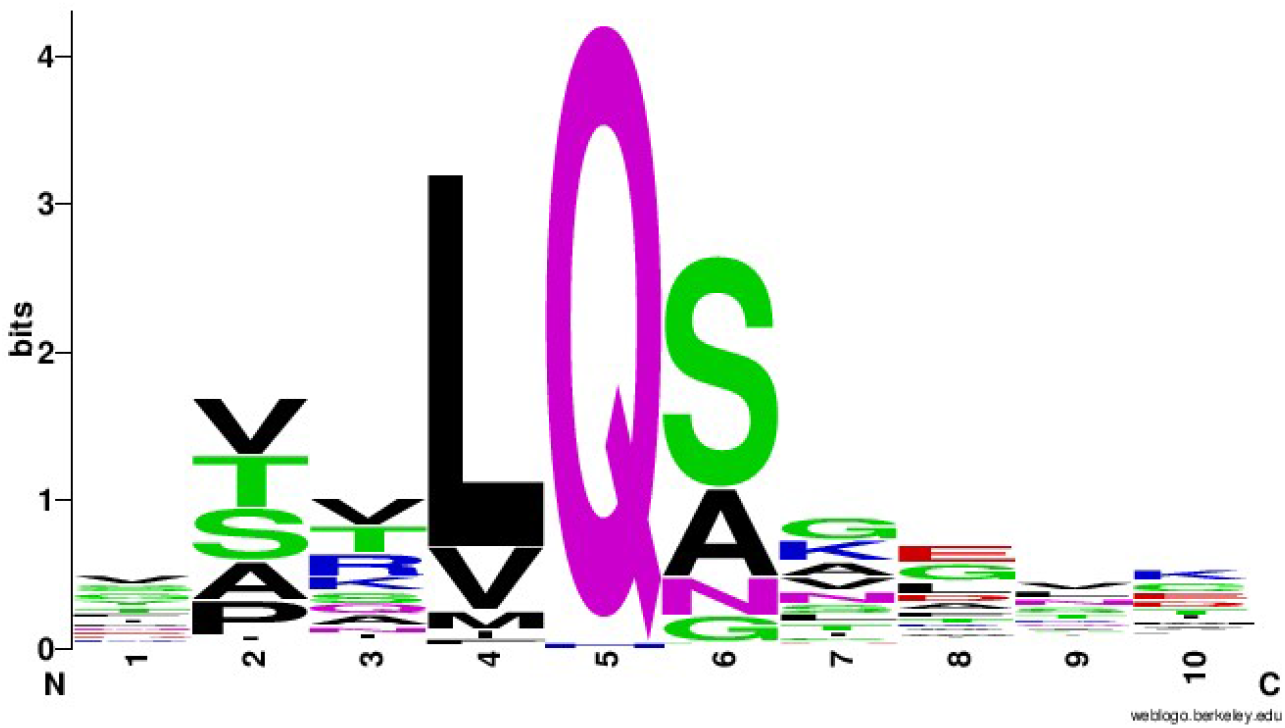
Improved 3CLpro cleavage site sequence logo plotted by WebLogo v2.8.2.[82]

**Figure 4:**
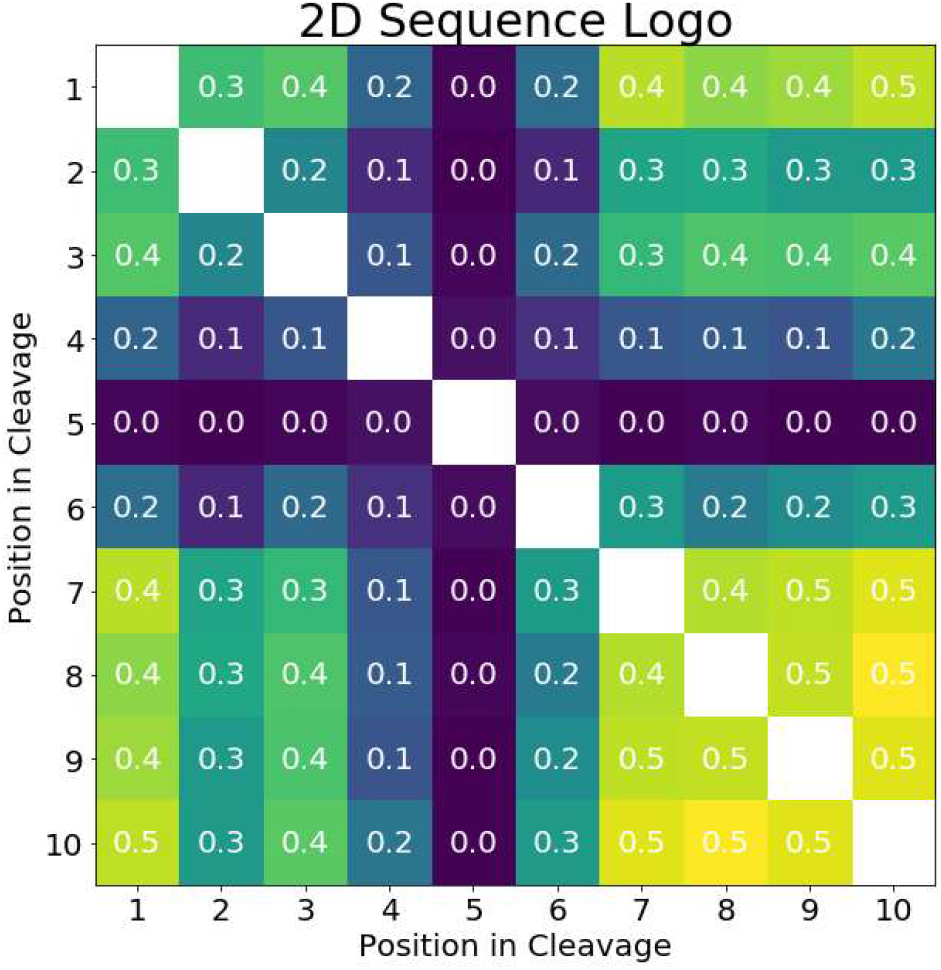
Entropy correlation coefficients (also known as symmetric uncertainties) between positions within the improved sequence logo.

**Figure 5:**
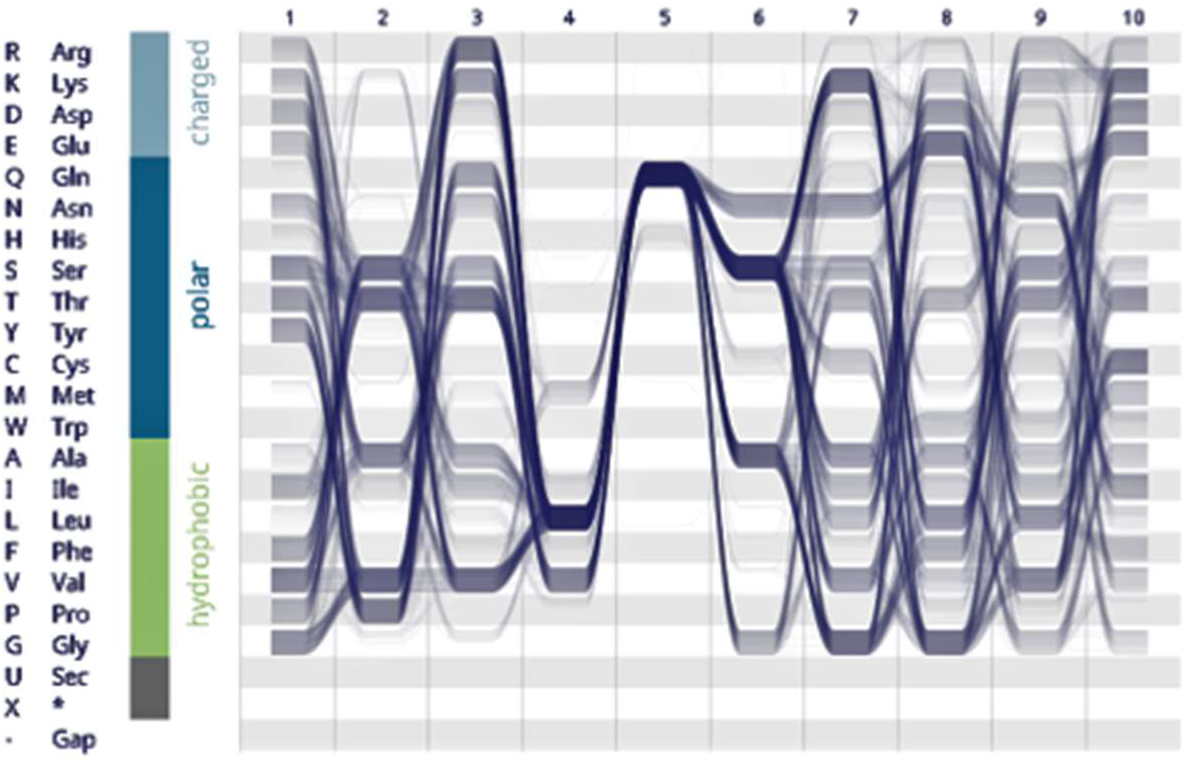
Sequence bundle with charge-polarity-hydrophobicity encoding.[83]

As for my improvements to the NN, note that Kiemer et al.’s MCC of 0.840 is an average from triple cross-validation (CV).[59] Because the known cleavage dataset is small, no data went unused; the three NN output scores were averaged and similarly considered cleavages when greater than 0.5. Applying this average scoring to the entire small and large dataset resulted in single MCCs of 0.946 and 0.849. Retraining the same NN structures (each with one hidden layer with 2 neurons) on the larger dataset resulted in three-average CV and single final MCCs of 0.979 and 0.996, a significant improvement even though the datasets are less balanced. Adding all other histidines (which precede 19/762 different cleavages) as negatives again improved the CV MCC to 0.994 and slightly reduced the final MCC to 0.992. Interestingly, two infectious bronchitis virus (IBV) polyproteins contained cleavages following leucine, methionine, and arginine (VSKLL^AGFKK in APY26744.1 and LVDYM^AGFKK and DAALR^NNELM in ADV71773.1). To my knowledge, synthetic tetra/octapeptides have been cleaved following histidine, phenylalanine, tryptophan, methionine, and possibly proline residues,[56, 84] but only one natural histidine substitution has been documented in HCoV-HKU1[85] and likely does not affect function.[86–88] To optimize hyperparameters, the whole dataset was repeatedly split into 80% training/20% testing sets with further splitting of the 80% training set for cross validation. The optimal settings, naive oversampling (within training folds[89]), averaged three-fold cross-validation (on the whole dataset, not just the initial 80%), limited-memory Broyden-Fletcher-Goldfarb-Shanno (lbfgs) solver, hyperbolic tangent activation, 0.00001 regularization, and 1 hidden layer with 10 neurons, had a 20% test set MCC average and standard deviation of 0.983+/−0.003 when split and trained many times. Train/test sets repeatedly split with different ratios in Figure 6 demonstrate that the entire dataset is not required for acceptable performance for all three classification methods, although the optimal and my finally method used three networks on all the data (with three-fold cross-validation), returning a three-average CV and final MCC of 0.983 and 0.998, respectively. Note that Figure 6 displays a curve for an equivalent physiochemical encoding (with input side 40 containing normalized volumes, interface and octanol hydrophobicity scales, and isoelectric points) underperforming when compared to one-hot encoding even at relatively small training sizes. Of these four physiochemical scales, octanol hydrophobicity alone reached a test MCC of 0.959, and, in the order of importance, addition of volume, interface hydrophobicity, and isoelectric point features increased the maximum test MCC to 0.977. Table 1 finally lists the one-hot encoded NN’s few incorrectly labeled sequences and their respective sources and scores.

**Figure 6:**
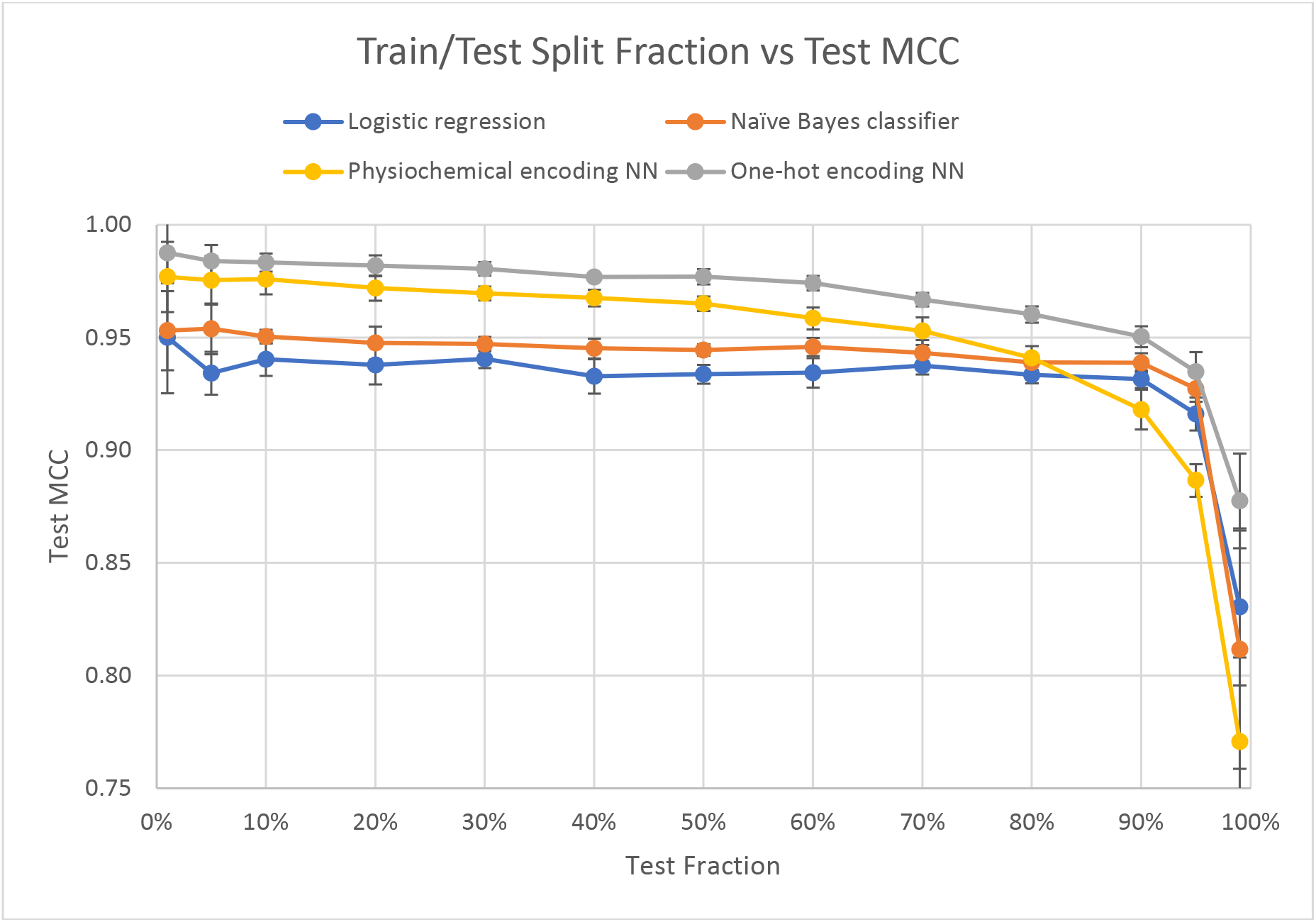
Train/test split fraction vs MCC demonstrating that performance quickly approaches a limit for all classifiers.

**Table 1:**
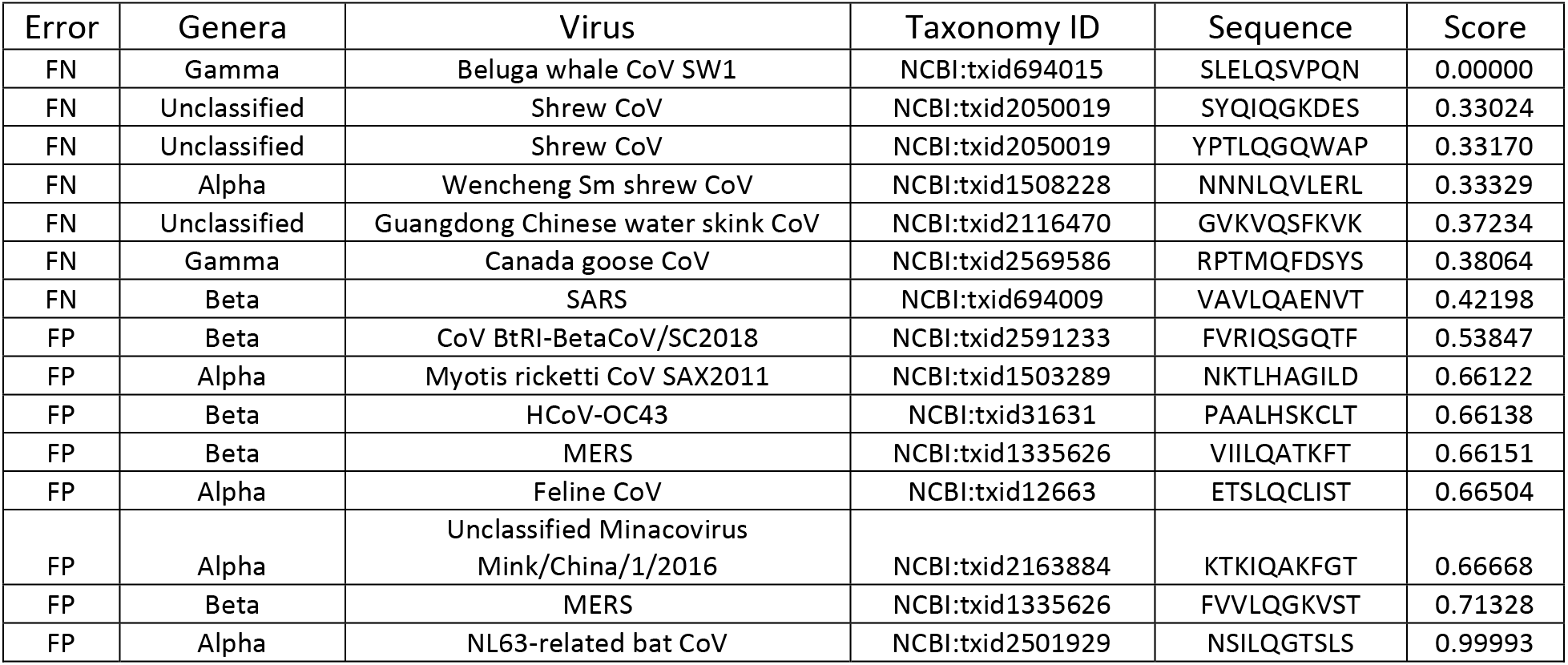
Only 15 out of 33,182 sequences were incorrectly labeled by the final NN. FN, false negative; FP, false positive.

Of the 20,350 manually reviewed human proteins, 4,460 were cleaved at least once with a NN score greater than or equal to 0.5. To prove that the 5,887 cleavages were nonrandomly distributed among human proteins (with a maximum of 25 cleavages in the 5,795 amino acid, RNA splicing regulation nucleoprotein protein AHNAK2), random sequences with weighted amino acid frequencies were checked for cleavages. Cleavages occurred at 1.10% of glutamines (4.77% of amino acids)[90] or every 1,900 amino acids in these random sequences. Most proteins are shorter than this and would, if randomly distributed, follow a Poisson distribution; my data’s deviation from this distribution indicates that many cleavages are intentional.

Tissue (UP_TISSUE and UNIGENE_EST_QUARTILE), InterPro, direct Gene Ontology (GO includes cellular compartment (CC), biological process (BP), and molecular function (MF)), Reactome pathways, sequence features, and keywords annotations were all explored in DAVID.[75, 76] Only annotations with Benjamini-Hochberg-corrected p-values less than 0.05 were considered statistically significant, and both enriched and depleted (no cleavages) annotations are listed in Tables S1–S9.

## Discussion

Enrichment and depletion analyses are often used to probe the importance of annotations in many disease states, yet quantification is not possible without experimentation. First, if a protein is central to a pathway, a single cleavage may be all that is required to generate equivalent downstream outcomes. Cleaved proproteins such as coagulation factors or complement proteins may even be activated by 3CLpro cleavage. Additional exhaustive analysis or inclusion of some measure of centrality is required to determine if any insignificantly enriched or depleted pathways are still affected at central nodes (i.e. false negatives). Second, protease-,[56] substrate sequence-,[77, 84, 91–93] substrate truncation-,[94] pH-, temperature-, inhibitor type and concentration-, and time after infection-dependent cleavage kinetics convert this classification problem into a regression problem. Cleavage rates among the 11 cleavages per pp1ab vary by at least 50-fold, so predictions here assume that 3CLpro exists in high enough concentrations and for a long enough time that rate constants do not matter because cleavage reactions are complete. Third, longer proteins are more likely to be randomly cleaved and may confound conclusions about annotations containing them. Cleavages in longer proteins (e.g. cytoskeletal or cell-cell adhesion components) are no less important than those in shorter sequences, and annotations containing proteins with multiple cleavages deviating from Poisson distributions are more likely due to highly conserved sequences than simply protein length. Lastly, convergent evolution within the host may also result in false positives and may be partially avoided by investigating correlations between domains, motifs, repeats, compositionally biased regions, or other sequence or structural similarities and other functional and ontological annotations. Ideally, a negative control proteome from an uninfectable species could prevent false positives, but coronaviruses are extremely zoonotic. Here, depletions in the human proteome are taken to be negative controls. Comparison with a bat proteome with deficiencies in many immune pathways, however, may show which human cleavages are unintentional or exerted little or no selective pressure before cross-species transmission.

### Tissues

As expected in this data, the most significant tissue enrichment of 3CLpro cleavages are in the epithelium, but central and peripheral nervous tissues are also affected due to their similar expression and enrichment of complex structural and cell junction proteins. It is noteworthy that major proteins associated with neurodegenerative disease are also predicted to be cleaved: Alzheimer’s disease (amyloid precursor protein (APP), tau protein), Parkinson’s disease (vacuolar protein sorting-associated VPS35, eukaryotic translation initiation factor EIF4G1, DNAJ homolog DNAJC13), Huntington’s disease (huntingtin), amyotrophic lateral sclerosis (trans-activation response element (TAR) DNA-binding TARDBP), and spinocerebellar ataxia type 1 (ataxin-1). Testis has somewhat similar expression to epithelium and brain, highly expresses ACE2, and is enriched in movement/motility-(subset of structural proteins) and meiosis-related (chromosome segregation) proteins, further increasing the likelihood that this tissue is infectible. Spleen, however, does not express much ACE2, and its enrichment is likely due to genes with immune function and mutagenesis sites. Proteins with greater tissue specificity (3^rd^ quartile) show additional enrichments along the respiratory tract (tongue, pharynx, larynx, and trachea), in immune tissues (lymph node and thymus), and in other sensory tissues (eye and ear). Combining tissues, tobacco use disorder is the only significantly enriched disease, but acquired immunodeficiency syndrome (AIDS) and atherosclerosis were surprisingly depleted.

Cleavages are also surprisingly depleted in olfactory and gustatory pathways given the virus’ ability to infect related cells and present as anosmia and ageusia. Olfactory receptors are transmembrane rhodopsin-like G protein-coupled receptors that, when bound to an odorant, stimulate production of cyclic adenosine monophosphate (cAMP) via the G protein and adenylate cyclase. The G proteins GNAL and GNAS are not cleaved, and some but not all adenylate cyclases are cleaved, likely resulting in an increase in cAMP. cAMP is mainly used in these cells to open their respective ligand-gated ion channels and cause depolarization, but it is also known to inhibit inflammatory responses through protein kinase A (PKA) and exchange factor directly activated by cAMP (EPAC). Multiple phosphodiesterases (PDEs) that degrade cAMP but not PDE4, the major PDE in inflammatory and immune cells, are cleaved. PDE4 inhibitors have been shown to reduce destructive respiratory syncytial virus-induces inflammation in lung,[95] but olfactory receptor neurons are quickly regenerated and sacrifice themselves when infected by influenza A virus.[96] The depletion in cleavages and resulting increase in cAMP in these neurons is likely to inhibit their programmed cell death long enough for the virus to be transmitted through the glomeruli to mitral cells and the rest of the olfactory bulb. Tongue infection may have similar mechanisms, and herpes simplex virus has been shown to be transmitted to the brainstem through the facial and trigeminal nerves.[97]

### Gene Ontology

Cleaved proteins are depleted in the extracellular space (except for structural collagen, laminin, and fibronectin mainly associated) and enriched in the cytoplasm and many of its components, indicating that the selective pressure for cleavage is weaker once cells are lysed and the protease is released. In the cytoplasm, the most obviously enriched sets are in the cytoskeleton (microfilament, intermediate filament, microtubule, and spectrin), motor proteins (myosin, kinesin, and dynein), cell adhesion molecules (integrin, immunoglobulin, cadherin, and selectin), and relevant Ras GTPases (Rho, Rab, Ran, Rac, and Arf), particularly in microtubule organizing centers (MTOCs) including centrosomes, an organelle central to pathways in the cell cycle including sister chromatid segregation. More specifically, cleavage of the cilia-associated proteins nephrocystins 1/2/4/5/6 (NPHP1/2/4/5/6), Bardet-Biedl syndromes proteins 1/9/12 (BBS1/9/12), Alstrom syndrome 1 (ALMS1), coiled-coil and C2 domain-containing protein 2A (CC2D2A), retinitis pigmentosa 1 (RP1), protein fantom (RPGRIP1), tubby-related protein 1 (TULP1), polycystin 1/2, protein kintoun (DNAAF2), dynein axonemal heavy chain 5/11 (DNAH5/11) and intermediate chain 2 (DNAI2), radial spoke head protein 6 homolog A (RSPH6A), and leucine-rich repeat-containing protein 50 (LRRC50) may contribute to dyskinesia and reduced mucociliary escalator effectiveness associated with many respiratory viruses including HCoV-229E and SARS and their resulting bacterial pneumonias.[98, 99] Additionally, cilial dysfunction in olfactory cells in COVID-19 leads to anosmia, although the main reported mechanism is nsp13 (helicase/triphosphatase)-centrosome interaction.[100] Coiled coils account for many of these cleavages and are primarily expressed in corresponding cellular compartments in the epithelium, testis, and brain. Only the coronavirus nsp1, nsp13, and spike proteins have so far been shown to interact with the cytoskeleton,[101–103] although many other viruses including influenza A virus,[104] herpes simplex virus, rabies virus, vesicular stomatitis virus, and adeno-associated virus[105] also modulate the cytoskeleton.[106] In neurons, this allows for axonal and trans-synaptic transport of viruses which can often be inhibited but sometimes exaggerated by cytoskeletal drugs often used in oncology.[107–110]

Modulation of these structural and motor proteins is required for formation of the double-membrane vesicles surrounding replicase complexes[111, 112] and for egress. Similarly required for vesicular transport, the coatomer COPI, clathrin, and caveolae pathways are untouched by 3CLpro other than the muscle-specific cavin-3, but COPII’s SEC24A/24B/31A are likely cleaved due to their function in selecting cargo[113, 114] and contribution to membrane curvature preventing inward nucleocapsid engulfment.[115] Cleavage of retromer component VSP35, ADP-ribosylation factor-binding protein GGA1, and many adaptor protein complexes (AP1B1/G1/G2, AP2A1/B1, AP3B1/B2/D1/M1/M2, and AP5B1/M1) often targeting degradation leaves only the poorly characterized AP4 or other unknown pathways to handle egress. Modulators of any of these vesicle trafficking pathways may be effective treatments for COVID-19.

The nucleus is enriched because its nuclear localization signals and scaffolding proteins are cleaved. Additionally, many nuclear pore complex proteins and importins/exportins associated with RNA transport are also cleaved. Lamins, which are cleaved by caspases during apoptosis to allow chromosome detachment and condensation, are also cleaved by 3CLpro. Chromatin-remodeling proteins including histone acetyltransferases (HATs) often containing bromodomains, histone deacetylases (HDACs), structural maintenance of chromosomes (SMC) proteins (cohesins and condensins) also containing coiled coils, separase (the cysteine protease that cleaves cohesin to separate sister chromatids), and topoisomerase III alpha, but not CCCTC-binding factor (CTCF) nor any other topoisomerases are cleaved, complicating the effects on chromosome condensation and global gene expression. HDAC inhibitors have been shown to decrease or increase virulence depending on the virus,[116–120] and some but not all DNA methyltransferases and demethylases are cleaved, further complicating these effects. Viruses benefit from preventing programmed cell death and its corresponding chromosomal compaction in response to viral infection (pyknosis), but they also attempt to reduce host transcription by condensing chromosomes and reroute translation machinery toward their own open reading frames.[121, 122] Relatedly, 28S rRNA has been shown to be cleaved by murine coronavirus, and ribosomes with altered activity are likely directed from host to viral RNAs.[123] Ribosome cleavages are depleted here because they are required for viral translation, but the few ribosomal proteins that are cleaved (RPL4/10 and RPS3A/19) tend to be more represented in monosomes, not polysomes,[124] indicating that ribosomes that initiate faster than they elongate are preferred because they likely frameshift more frequently, allowing for control of the stoichiometric ratio of pp1a and pp1ab.[125] If slower ribosomes are not directly more likely to frameshift, they are still less likely to participate if frameshift-induced traffic jams, collision-stimulated translation abortion and splitting,[126] and subsequent 60S subunit obstruction sensing and nascent-chain ubiquitylation, which is especially noteworthy because zinc finger 598 (ZNF598), nuclear export mediator factor (NEMF), and listerin E3 ubiquitin ligase 1 (LTN1) are predicted to be cleaved.[127] Signal recognition particle (SRP) subunits 54/68/72kDa associated with the ribosome are also predicted to be cleaved. SRP, especially the uncleaved SRP9/14kDa heterodimer, encourage translation elongation arrest to allow translocation including transmembrane domain insertion (e.g. coronavirus envelope protein) and has been associated with frameshifts.[128–130] In fact, frameshifting is a highly enriched keyword in cleaved proteins mainly due to endogenous retroviral (ERV) elements, some of which can activate an antiviral response via pattern recognition receptors (PRRs).[131] Some also resemble reverse transcriptases and may, like the CRIPSR system in prokaryotes, be capable of copying coronavirus genomic RNA to produce an RNAi response via the similarly cleaved DICER1, AGO1/2, and PIWL1/3.[132] If the latter is true, individuals with distinct ERV alleles and loci may differentially respond to SARS-CoV-2 infection and/or treatment, especially exogenous RNAi. Lastly, ribosomal proteins are also included in the nonsense-mediated decay (NMD) pathway, which is likely depleted in cleavages because NMD has been shown to be a host defense against coronavirus genomic and subgenomic RNAs’ multiple ORFs and large 3’ UTRs.[133] It was also shown that the nucleocapsid protein inhibits this degradation but often cannot protect newly synthesized RNAs early in infection. The selective pressure on 3CLpro may be reversed by this nucleocapsid inhibition and the preferential degradation of host mRNAs such that host resources can again be directed toward viral translation.

In addition to affecting large organelles, 3CLpro is predicted to cleave all known components of vault: major vault proteins (MVP), telomerase protein component 1 (TEP1), and poly(ADP-ribose) polymerase 4 (PARP4). Vault function has not been completely described, but it has known interactions with other viruses.[134–136] Telomerase reverse transcriptase (TERT) is also cleaved, but is more frequently reported to be activated by other viral infections and/or promote oncogenesis.[137]

Other common viral process proteins are enriched in the epithelium and adaptive immune cells, and those cleaved may affect the heat shock response and other small RNA processing. Lactoferrin, an antiviral protein that is upregulated in SARS infection,[138] is also cleaved, although one of its fragments, lactoferricin, has known antiviral activity.[139] Cleaved PRRs include the toll-like receptors TLR6/8; the C-type lectin receptors CLEC4G/H1/4K/4L/10A/13B/13C/16A; NK cell lectin-like receptors (KLRC4/G1), aspartate/glutamate carrier 1 (ACG1), collectin-7/12, neurocan core protein, FRAS1-related extracellular matrix protein 1 (FREM1), layilin, polycystin 1, E-selectin, and thrombomodulin primarily present on dendritic cells; and the NOD-like receptors NOD2 and NLRP1/2/3/6/10/12/14. Cleaved proteins downstream of these PRRs include receptor-interacting serine-threonine-protein kinase 1/2 (RIP1/2), NF-kappa-B p100 subunit, CASP8 and FADD-like apoptosis regulator (CFLAR), TIR-domain-containing adapter-inducing interferon-beta (TRIF), IRF2, and death domain-associated protein 6 (DAXX), and other relevant downstream pathways similarly include many cleaved proteins: phosphoinositide 3-kinase/protein kinase B (PI3K/AKT) pathway (PIK3CG/D, PIK3R2/5/6, serine/threonine-protein phosphatase 2A (PPP2R1A/2A/2B/2C/2D/3B/5B)(PP2A dephosphorylates AKT), neuronal and inducible nitric oxide synthase (n/iNOS)(where nitric oxide has conflicting effects on viral infections[140, 141]), tuberous sclerosis 1/2 (TSC1/2)); mechanistic target of rapamycin (mTOR) pathway (ribosomal S6 kinase 1/2, sterol regulatory element-binding protein 1 (SREBP1), RB1-inducible coiled-coil protein 1 (RBCC1), regulatory-associated protein of mTOR (RAPTOR), rapamycin-insensitive companion of mTOR (RICTOR), proline-rich 5-like (PRR5L), CAP-Gly domain containing linker protein 1 (CLIP1), forkhead box protein O1/3 (FOXO1/3)); and mitogen-activated protein kinase (MAPK) pathway (MAP4K2/4/5, MAP3K4/5/6/8/13/14, kinase suppressor of RAS 2 (KSR2), MAP2K3/4, MAPK7/13/15, dual specificity protein phosphatase 8 (DUSP8), and mouse double minute 2 homolog (MDM2)). Additional cleaved transcription factors include c-Jun, activating transcription factor 6 (ATF6), the cAMP-responsive element-binding proteins CREB3/5/BP, specificity protein 1 (SP1), octamer transcription factors OCT1/2, the heat schock factors HSF2/2BP/4/X1, RNA polymerase I initiator nucleolar transcription factor 1 (UBTF), RNA polymerase II initiators TFIID and selective factor 1 (SL1) subunits (TATA-binding protein (TBP), TBP-like 2, TBP-associated factor 1C/6/172) and mediator coactivator subunits 1/12L/13/15/17/22/26/28, RNA polymerase III initiators TFIIIB150, TFIIIC, and snRNA-activating protein complex subunit 4 (SNAPC4). No interferons are cleaved likely due to their redundancy, and no interferon receptors are cleaved. The downstream effectors STAT1/2/4; the ISGs guanylate-binding protein 1 (GBP1), interferon alpha-inducible protein 6 (IFI6), membrane spanning 4-domain A4A (MS4A4A), 2’-5’-oligoadenylate synthetase 1 (OAS1), promyelocytic leukemia protein (PML), mitoferrin-2, three prime repair exonuclease 1 (TREX1), and tripartite motif-containing protein 5 (TRIM5); and the tumor necrosis factor (TNF) ligands (TNFSF3/13/18 and ectogysplasin A) and receptor TNFRSF21 are, however, also cleaved. Finally, pro-apoptotic protein cleavages exist in the Bcl-2 family (Bcl-rambo) and in caspases (CASP2/5/12), although the anti-apoptotic Bcl-2 protein (Bcl-B) and inhibitors of apoptosis (baculoviral IAP repeat-containing proteins BIRC2/3/6) are also cleaved.

### Other Pathways and Keywords

Lipoproteins are a depleted keyword, but apolipoproteins APOA-V/B/L1/(a), cholesteryl ester transfer protein (CETP), microsomal triglyceride transfer protein (MTTP), and the lipid transfer receptors LDL-related proteins LRP2/6/12 are all predicted to be cleaved and, other than the proapoptotic APOL1,[142] are associated with chylomicrons, VLDL, and LDL as opposed to HDL, indicating that lipoproteins may contribute to the correlations between COVID-19 symptom severity, dyslipidemia, and cardiovascular disease. It was recently discovered that SARS-CoV-2 spike protein binds cholesterol, allowing for association with and reduced serum concentration of HDL. These findings combined with the 3CLpro cleavages show an opportunity for HDL receptor inhibitor treatment, especially antagonists of the uncleaved scavenger receptor SR-B1.[143] Cleavage of the adipokines leptin, leptin receptor, and IL-6 provide a mechanism for COVID-19 comorbidity with obesity independent of lipoproteins and indicate another potential treatment: anti-leptin antibodies.[144, 145]

Ubiquitinating and deubiquitinating (DUBs) enzymes are most enriched in the epithelium and the nucleus and include cleaved ubiquitin ligase-supporting scaffolding cullins and DUBs such as the ubiquitous proteasomal subunit RPN11 and related lid subunits RPN6/10/12. Ubiquitin itself is not, but neural precursor cell expressed developmentally downregulated protein 4 (NEDD4) and the related SMAD ubiquitination regulatory factor 1/2 (SMURF1/2) and HECT, C2, and WW domain containing E3 ubiquitin ligase 1 (HECW1) are, cleaved. NEDD4 has been shown to enhance influenza infectivity by inhibiting interferon-induced transmembrane protein 3 (IFITM3)[146, 147] and Japanese encephalitis virus by inhibiting autophagy,[148] but its ubiquitination of many diverse human viruses promotes their egress. IFITMs generally have antiviral activity (others include HIV-1,[149] dengue virus,[150] and filoviruses[151]), but its use as a treatment for COVID-19 should be carefully considered given its varying effects among other coronaviruses.[152, 153] SARS-CoV-2 has two probable NEDD4 binding sites: the proline-rich, N-terminal PPAY and LPSY[154] in the spike protein and nsp8, respectively. Although the former sequence is APNY and is likely not ubiquitinated in SARS-CoV, small molecule drugs targeting this interaction or related kinases may be useful treatments for COVID-19 as they have been for other RNA viruses.[155–157] Further research is required to compare these cleavages to the PLpro deubiquitinating activity and the specificity and function of distinct ubiquitin and other ubiquitin-like protein linkage sites.[158, 159]

Helicases make up approximately 1% of eukaryotic genes and are enriched in cleavages with many containing RNA-specific DEAD/DEAH boxes. Most viruses except for retroviruses have their own helicase (nsp13 in SARS-CoV-2) and multiple human RNA helicases have been shown to sense viral RNA or enhance viral replication.[160–162] SARS nsp13 and nsp14 have been shown to be enhanced by the uncleaved human DDX5 and DDX1, respectively.[163, 164], however multiple proteins interacting with DDX1 (FAM98A) and DDX5 (DHX15, SNW1, MTREX, and HNRNPH1), the retinoic acid-inducible gene I (RIG-I)-associating DDX6, and DDX20 involved in ribosome assembly are predicted to be cleaved, making these effects enigmatic without knowledge of additional interactions with other nsps.

Fibronectin type-III domains are enriched, but fibronectin itself is not cleaved. No cleaved proteins with this domain are directly related to coagulation, but the related tissue factor, coagulation factors VIII (antihemophilic factor A glycoprotein, also an acute-phase protein secreted in response to infection), XII (Hageman factor), XIII (fibrin-stabilizing factor transglutaminase), plasmin(ogen), von Willebrand factor, and kininogen-1 are cleaved. Multiple cleaved serpin suicide protease inhibitors (plasminogen activator inhibitor-2, megsin, alpha-1-antitrypsin, and the less relevant angiotensinogen, protein Z-dependent protease inhibitor, leukocyte elastase inhibitor, and heat shock protein 47) are also related to coagulation, potentially increasing both thrombosis and fibrinolysis rates or resulting in dose-dependent effects.[165, 166] Angiotensinogen is, however, unrelated to coagulation and is cleaved far from its N-terminus, so its effects on the renin-angiotensin system remain unknown. The structurally similar alpha-2-macroglobulin has a predicted cleavage outside its protease bait region, however, the addition of a missense mutation Q694S would allow cleavage at the same site as factor XIII without reducing protease trapping ability as much as large deletions.[167, 168] Additional support for this potential exogenous replacement includes presence of serine in the same position in pregnancy zone protein (PZP), which shares 71% identity with alpha-2-macroglobin and contains a neighboring GAG site resembling known PLpro cleavages in its primary bait region. Most other antiproteases, however, are too small to have many potential cleavage sites even though they are a very important response to respiratory virus infection. Serpin or alpha globulin replacement therapy or treatment with modified small, 3CLpro competitive inhibitors may be a useful treatment for COVID-19.[169]

In addition to coagulation factors, the complement system can induce, in addition to many other components of innate immunity, expulsion of neutrophil extracellular traps (NETs) intended to bind and kill pathogens.[170] NETs, however, simultaneously trap platelets expressing tissue factor and contribute to hypercoagulability. The complement pathway is not obviously enriched, but many central proteins (C1/3/4 and factor B) are or have subunits that are cleaved, indicating viral adaptation to the classical, alternative, and likely lectin pathways.[171–173] Neutrophilia and NET-associated host damage are known to occur in severe SARS-CoV-2 infection, so inhibitors of the pathway are currently in clinical trials: histone citrullination, neutrophil elastase, and gasdermin D inhibitors to prevent release and DNases to degrade chromatin after release.[174, 175] Complement inhibition would likely similarly reduce the risks of hypercoagulability and other immune-mediated inflammation associated with COVID-19, but effects may vary widely between sexes and ages.[176, 177]

Redox-active centers including proteins involved in selenocysteine synthesis are additionally depleted in cleavages likely because of their involvement in avoiding cell death and innate immune response. Respiratory viruses differentially modulate redox pathways, balancing lysis-enhanced virion proliferation and dual oxidase 2 (DUOX2)-derived reactive oxygen species (ROS)-induced interferon response.[178] In addition to depleted antioxidant proteins, cleavage of DUOX1/2, NADPH oxidase 5 (NOX5), and xanthine oxidase (XO), the former of which are upregulated in chronic obstructive pulmonary disease (COPD),[179] indicates that coronaviruses prefer to reduce oxidative stress in infected cells, contrary to most COVID-19 symptoms. Given the diversity of responses to respiratory virus infections, each proposed antioxidant should be thoroughly evaluated before being recommended as a treatment of COVID-19.

The impact of post-translational modifications on viral protease cleavage frequency remains uncharacterized. Glutamine and leucine, the two most important residues in the cleavage sequence logo, are rarely modified, but serine, the next most important residue, is the most frequently phosphorylated amino acid. Analysis of keywords showed enrichment of phosphoproteins and depletion of disulfide crosslinked, lipid-anchored, and other transmembrane proteins.

Lastly, the keywords polymorphism and alternate splicing were enriched, indicating that additional variability between cell lines and between individuals are likely. Once health systems are not so burdened by the quantity of cases and multiple treatments are developed, personalized interventions will likely differ significantly between individuals.

## Conclusion

Many expected and novel protein annotations were discovered to be enriched and depleted in cleavages, indicating that 3CLpro is a much more important virulence factor than previously believed. 3CLpro cleavages are enriched in the epithelium (especially along the respiratory tract), brain, testis, plasma, and immune tissues and depleted in olfactory and gustatory receptors. Affected pathways with discussed connections to viral infections include cytoskeleton/motor/cell adhesion proteins, nuclear condensation and other epigenetics, host transcription and RNAi, coagulation, pattern recognition receptors, growth factor, lipoprotein, redox, ubiquitination, apoptosis. These pathways point toward many potential therapeutic mechanisms to combat COVID-19: cytoskeletal drugs frequently used against cancer, modulators of ribosomal stoichiometry to enrich monosomes, upregulation of DICER1 and AGO1/2, exogenous lactoferrin and modified antiproteases including alpha globulins, upregulation of serpins potentially via dietary antioxidants, complement inhibition, reduction of LDL and inhibition of HDL receptor (e.g. by antagonizing SR-B1), anti-leptin antibodies, and downregulating NEDD4 or related kinases and upregulating IFITMs. Pathway components with more complex disruption that may also deliver therapeutic targets but require elucidating experimental results include PDEs, histone acetylation, nitric oxide, and vesicle coatomers. It is also worth further investigating how 3CLpro contributes if at all to the correlations between obesity and severity of infection or to viral induction of autoimmune and potentially oncological conditions.

Expansion of the training dataset to the whole order *Nidovirales* may provide more diversity to improve classifying methods if additional protease/cleavage coevolution does not invalidate the assumption of cross-reactivity. Issues requiring *in vitro* and *in vivo* experimentation include characterization of cleavage kinetics, any functional differences between proteases, the molecular effects of post-translation modifications, the individual and population effects of polymorphisms in cleavage sequences on susceptibility to or severity of infection. Even though many caveats exist without experimentation, similar prediction, enrichment/depletion analysis, and therapeutic target identification should be performed for every other viral protease.

## Acknowledgments

I am very grateful for my mother, Victoria Prescott, Esq., and friends who have given me invaluable help and advice throughout my work on this project.

## Supplementary Information

**Table S1:** Significant UP_TISSUE enrichments and depletions.

| Enriched |  |  |  | Depleted |  |  |  |
| --- | --- | --- | --- | --- | --- | --- | --- |
| Tissue | FE | p-value | Benjamini | Tissue | FE | p-value | Benjamini |
| Plasma | 1.56 | 1.40E-06 | 1.64E-04 | Hair root | 1.30 | 1.22E-05 | 3.95E-03 |
| Fetal kidney | 1.55 | 1.58E-04 | 8.14E-03 | Umbilical cord blood | 1.17 | 6.56E-09 | 4.24E-06 |
| Hepatoma | 1.50 | 8.76E-05 | 5.83E-03 | Cajal-Retzius cell | 1.16 | 1.69E-05 | 3.63E-03 |
| Epithelium | 1.44 | 1.50E-37 | 7.01E-35 |  |  |  |  |
| Amygdala | 1.32 | 3.39E-05 | 2.63E-03 |  |  |  |  |
| Teratocarcinoma | 1.31 | 1.62E-04 | 7.55E-03 |  |  |  |  |
| Spleen | 1.26 | 3.12E-05 | 2.91E-03 |  |  |  |  |
| Testis | 1.21 | 1.22E-17 | 1.90E-15 |  |  |  |  |
| Brain | 1.18 | 1.80E-30 | 4.21E-28 |  |  |  |  |

**Table S2:** Significant UNIGENE_EST_QUARTILE enrichments and depletions.

| Enriched |  |  |  | Depleted |  |  |  |
| --- | --- | --- | --- | --- | --- | --- | --- |
| Tissue | FE | p-value | Benjamini | Tissue | FE | p-value | Benjamini |
| larynx_normal_3rd | 1.35 | 1.50E-26 | 1.14E-24 | salivary gland_normal_3rd | 1.05 | 2.77E-06 | 2.10E-04 |
| oral tumor_disease_3rd | 1.35 | 8.45E-24 | 1.60E-22 | neonate (< 4 weeks old)_development_3rd | 1.03 | 7.94E-04 | 1.50E-02 |
| pharynx_normal_3rd | 1.33 | 2.62E-14 | 1.99E-13 | non-glioma_disease_3rd | 1.03 | 1.49E-04 | 5.65E-03 |
| laryngeal cancer_disease_3rd | 1.33 | 2.42E-24 | 6.14E-23 | bone marrow_normal_3rd | 1.03 | 5.42E-04 | 1.36E-02 |
| tongue_normal_3rd | 1.31 | 6.07E-26 | 2.31E-24 | heart_normal_3rd | 1.02 | 9.88E-04 | 1.49E-02 |
| thyroid_normal_3rd | 1.26 | 3.36E-19 | 5.11E-18 | skin_normal_3rd | 1.02 | 1.59E-03 | 1.99E-02 |
| trachea_normal_3rd | 1.26 | 3.56E-16 | 3.66E-15 |  |  |  |  |
| pharyngeal tumor_disease_3rd | 1.25 | 1.93E-09 | 1.22E-08 |  |  |  |  |
| thyroid tumor_disease_3rd | 1.23 | 1.11E-14 | 9.37E-14 |  |  |  |  |
| mammary gland_normal_3rd | 1.22 | 8.77E-18 | 1.11E-16 |  |  |  |  |
| colorectal tumor_disease_3rd | 1.22 | 1.07E-14 | 1.01E-13 |  |  |  |  |
| breast (mammary gland) cancer_disease_3rd | 1.19 | 3.06E-11 | 2.11E-10 |  |  |  |  |
| adipose tissue_normal_3rd | 1.16 | 4.51E-07 | 2.28E-06 |  |  |  |  |
| colon_normal_3rd | 1.15 | 2.51E-09 | 1.47E-08 |  |  |  |  |
| uterine tumor_disease_3rd | 1.13 | 5.07E-07 | 2.41E-06 |  |  |  |  |
| eye_normal_3rd | 1.13 | 2.10E-08 | 1.14E-07 |  |  |  |  |
| muscle_normal_3rd | 1.12 | 7.95E-06 | 3.36E-05 |  |  |  |  |
| lymph node_normal_3rd | 1.12 | 4.02E-06 | 1.80E-05 |  |  |  |  |
| thymus_normal_3rd | 1.11 | 2.26E-04 | 9.02E-04 |  |  |  |  |
| ear_normal_3rd | 1.09 | 6.52E-03 | 2.05E-02 |  |  |  |  |
| pituitary gland_normal_3rd | 1.09 | 4.07E-03 | 1.34E-02 |  |  |  |  |
| connective tissue_normal_3rd | 1.08 | 8.06E-03 | 2.43E-02 |  |  |  |  |
| chondrosarcoma_disease_3rd | 1.08 | 1.63E-03 | 5.90E-03 |  |  |  |  |
| testis_normal_3rd | 1.07 | 6.92E-04 | 2.63E-03 |  |  |  |  |

**Table S3:** Significant InterPro enrichments and depletions.

| Enriched |  |  |  | Depleted |  |  |  |
| --- | --- | --- | --- | --- | --- | --- | --- |
| Pfam | FE | p-value | Benjamini | Pfam | FE | p-value | Benjamini |
| Dynein heavy chain | 4.50 | 1.95E-09 | 6.57E-07 | High sulphur keratin-associated protein | 1.29 | 1.64E-05 | 1.81E-02 |
| Dynein heavy chain domain | 4.50 | 1.95E-09 | 6.57E-07 | Small GTP-binding protein domain | 1.21 | 1.20E-07 | 2.35E-04 |
| Dynein heavy chain, coiled coil stalk | 4.50 | 1.95E-09 | 6.57E-07 | Thioredoxin-like fold | 1.20 | 5.50E-06 | 7.13E-03 |
| Dynein heavy chain, domain-2 | 4.50 | 1.95E-09 | 6.57E-07 | Olfactory receptor | 1.20 | 1.35E-17 | 3.51E-14 |
| Dynein heavy chain, P-loop containing D4 domain | 4.50 | 8.26E-09 | 2.35E-06 | GPCR, rhodopsin-like, 7TM | 1.18 | 1.72E-22 | 1.35E-18 |
| ATPase, dynein-related, AAA domain | 4.50 | 3.48E-08 | 7.60E-06 | G protein-coupled receptor, rhodopsin-like | 1.17 | 8.42E-22 | 3.29E-18 |
| Peptidase A2A, retrovirus RVP subgroup | 4.50 | 1.04E-05 | 9.89E-04 | Kruppel-associated box | 1.13 | 6.74E-07 | 1.05E-03 |
| Dynein heavy chain, domain-1 | 4.50 | 4.24E-05 | 3.41E-03 |  |  |  |  |
| Retroviral nucleocapsid protein Gag | 4.50 | 4.24E-05 | 3.41E-03 |  |  |  |  |
| Beta-retroviral matrix, N-terminal | 4.50 | 4.24E-05 | 3.41E-03 |  |  |  |  |
| PH-BEACH domain | 4.50 | 1.71E-04 | 1.08E-02 |  |  |  |  |
| Na/K/Cl co-transporter superfamily | 4.50 | 6.77E-04 | 3.62E-02 |  |  |  |  |
| Peptidase A2A, retrovirus, catalytic | 4.09 | 4.59E-05 | 3.54E-03 |  |  |  |  |
| Retrovirus capsid, N-terminal core | 4.05 | 1.70E-04 | 1.10E-02 |  |  |  |  |
| Myosin-like IQ motif-containing domain | 4.03 | 1.64E-08 | 4.33E-06 |  |  |  |  |
| BEACH domain | 4.00 | 6.20E-04 | 3.37E-02 |  |  |  |  |
| Retroviral envelope protein | 4.00 | 6.20E-04 | 3.37E-02 |  |  |  |  |
| MyTH4 domain | 4.00 | 6.20E-04 | 3.37E-02 |  |  |  |  |
| Myosin, N-terminal, SH3-like | 3.90 | 3.24E-06 | 3.54E-04 |  |  |  |  |
| Myosin tail | 3.79 | 2.35E-07 | 3.63E-05 |  |  |  |  |
| Spectrin/alpha-actinin | 3.57 | 1.18E-09 | 4.38E-07 |  |  |  |  |
| Spectrin repeat | 3.42 | 1.55E-07 | 2.61E-05 |  |  |  |  |
| Laminin, N-terminal | 3.38 | 9.01E-05 | 6.53E-03 |  |  |  |  |
| Myosin head, motor domain | 3.04 | 7.82E-09 | 2.42E-06 |  |  |  |  |
| Mitochondrial carrier protein | 3.00 | 6.15E-06 | 6.16E-04 |  |  |  |  |
| Peptidase aspartic, active site | 3.00 | 1.02E-04 | 6.99E-03 |  |  |  |  |
| G-patch domain | 2.96 | 1.09E-06 | 1.39E-04 |  |  |  |  |
| SNF2-related | 2.96 | 1.09E-06 | 1.39E-04 |  |  |  |  |
| Actinin-type, actin-binding, conserved site | 2.94 | 7.14E-05 | 5.28E-03 |  |  |  |  |
| Cadherin, N-terminal | 2.91 | 1.63E-12 | 6.71E-10 |  |  |  |  |
| Arf GTPase activating protein | 2.85 | 8.68E-06 | 8.46E-04 |  |  |  |  |
| IQ motif, EF-hand binding site | 2.85 | 1.34E-15 | 9.88E-13 |  |  |  |  |
| Mitochondrial carrier domain | 2.63 | 5.71E-08 | 1.11E-05 |  |  |  |  |
| Mitochondrial substrate/solute carrier | 2.63 | 5.71E-08 | 1.11E-05 |  |  |  |  |
| Bromodomain, conserved site | 2.60 | 4.26E-04 | 2.40E-02 |  |  |  |  |
| Cadherin conserved site | 2.59 | 1.97E-15 | 1.23E-12 |  |  |  |  |
| HECT | 2.57 | 2.86E-04 | 1.67E-02 |  |  |  |  |
| Cadherin | 2.52 | 7.39E-15 | 3.45E-12 |  |  |  |  |
| Cadherin-like | 2.51 | 5.04E-15 | 2.65E-12 |  |  |  |  |
| Aspartic peptidase | 2.50 | 7.04E-04 | 3.66E-02 |  |  |  |  |
| Laminin G domain | 2.48 | 1.38E-07 | 2.44E-05 |  |  |  |  |
| Forkhead-associated (FHA) domain | 2.43 | 9.60E-05 | 6.69E-03 |  |  |  |  |
| Kinesin, motor region, conserved site | 2.42 | 4.36E-05 | 3.43E-03 |  |  |  |  |
| Dbl homology (DH) domain | 2.41 | 3.34E-08 | 7.75E-06 |  |  |  |  |
| Bromodomain | 2.41 | 2.94E-05 | 2.48E-03 |  |  |  |  |
| Kinesin, motor domain | 2.40 | 1.99E-05 | 1.71E-03 |  |  |  |  |
| Armadillo-like helical | 2.38 | 1.70E-23 | 3.15E-20 |  |  |  |  |
| Calponin homology domain | 2.31 | 9.23E-08 | 1.71E-05 |  |  |  |  |
| Ubiquitin-associated/translation elongation factor EF1B, N-terminal, eukaryote | 2.25 | 1.62E-04 | 1.07E-02 |  |  |  |  |
| GPS domain | 2.25 | 7.84E-04 | 3.90E-02 |  |  |  |  |
| Rho GTPase activation protein | 2.23 | 3.32E-08 | 8.21E-06 |  |  |  |  |
| EGF-like, laminin | 2.19 | 7.78E-04 | 3.93E-02 |  |  |  |  |
| Zinc finger, PHD-finger | 2.17 | 1.09E-06 | 1.44E-04 |  |  |  |  |
| Rho GTPase-activating protein domain | 2.15 | 1.08E-05 | 1.00E-03 |  |  |  |  |
| Band 4.1 domain | 2.12 | 2.33E-04 | 1.45E-02 |  |  |  |  |
| FERM domain | 2.12 | 2.33E-04 | 1.45E-02 |  |  |  |  |
| FERM central domain | 2.12 | 2.33E-04 | 1.45E-02 |  |  |  |  |
| Armadillo-type fold | 2.09 | 7.25E-24 | 2.69E-20 |  |  |  |  |
| Pleckstrin homology domain | 2.04 | 2.71E-17 | 2.51E-14 |  |  |  |  |
| WW domain | 2.04 | 4.62E-04 | 2.56E-02 |  |  |  |  |
| Zinc finger, PHD-type | 2.02 | 4.39E-06 | 4.65E-04 |  |  |  |  |
| Zinc finger, PHD-type, conserved site | 2.02 | 9.39E-05 | 6.67E-03 |  |  |  |  |
| Zinc finger, FYVE/PHD-type | 1.95 | 4.92E-08 | 1.01E-05 |  |  |  |  |
| Helicase, superfamily 1/2, ATP-binding domain | 1.92 | 3.08E-06 | 3.45E-04 |  |  |  |  |
| Collagen triple helix repeat | 1.92 | 7.03E-05 | 5.31E-03 |  |  |  |  |
| Pleckstrin homology-like domain | 1.92 | 2.16E-21 | 2.67E-18 |  |  |  |  |
| Intermediate filament protein, conserved site | 1.90 | 7.00E-04 | 3.69E-02 |  |  |  |  |
| Helicase, C-terminal | 1.88 | 1.12E-05 | 1.01E-03 |  |  |  |  |
| Peptidase C19, ubiquitin carboxyl-terminal hydrolase 2 | 1.86 | 2.52E-04 | 1.54E-02 |  |  |  |  |
| AAA+ ATPase domain | 1.85 | 1.25E-06 | 1.55E-04 |  |  |  |  |
| PDZ domain | 1.83 | 4.30E-07 | 6.14E-05 |  |  |  |  |
| Immunoglobulin I-set | 1.82 | 1.99E-06 | 2.38E-04 |  |  |  |  |
| Galactose-binding domain-like | 1.81 | 4.25E-04 | 2.43E-02 |  |  |  |  |
| Intermediate filament protein | 1.79 | 8.40E-04 | 4.12E-02 |  |  |  |  |
| von Willebrand factor, type A | 1.76 | 2.70E-04 | 1.60E-02 |  |  |  |  |
| Immunoglobulin E-set | 1.73 | 2.56E-04 | 1.54E-02 |  |  |  |  |
| Src homology-3 domain | 1.71 | 1.58E-07 | 2.55E-05 |  |  |  |  |
| Fibronectin, type III | 1.63 | 5.57E-06 | 5.73E-04 |  |  |  |  |
| Ankyrin repeat-containing domain | 1.60 | 9.58E-07 | 1.32E-04 |  |  |  |  |
| Ankyrin repeat | 1.59 | 2.11E-06 | 2.44E-04 |  |  |  |  |
| Epidermal growth factor-like domain | 1.56 | 1.78E-05 | 1.57E-03 |  |  |  |  |
| Immunoglobulin subtype 2 | 1.51 | 3.77E-05 | 3.10E-03 |  |  |  |  |
| EGF-like, conserved site | 1.47 | 7.18E-04 | 3.68E-02 |  |  |  |  |
| P-loop containing nucleoside triphosphate hydrolase | 1.33 | 2.45E-07 | 3.64E-05 |  |  |  |  |
| Immunoglobulin subtype | 1.32 | 1.42E-04 | 9.53E-03 |  |  |  |  |

**Table S4:** Significant GO CC enrichments and depletions.

| Enriched |  |  |  | Depleted |  |  |  |
| --- | --- | --- | --- | --- | --- | --- | --- |
| CC | FE | p-value | Benjamini | CC | FE | p-value | Benjamini |
| spectrin | 4.55 | 3.94E-05 | 3.16E-03 | ribosome | 1.17 | 2.45E-05 | 1.76E-03 |
| axonemal dynein complex | 4.09 | 1.58E-04 | 7.50E-03 | mitochondrion | 1.05 | 8.74E-05 | 4.13E-02 |
| spectrin-associated cytoskeleton | 3.98 | 2.08E-03 | 4.83E-02 | integral component of membrane | 1.03 | 3.75E-08 | 5.43E-05 |
| nuclear pore nuclear basket | 3.79 | 1.36E-04 | 7.44E-03 |  |  |  |  |
| microtubule plus-end | 3.74 | 2.65E-06 | 4.62E-04 |  |  |  |  |
| myosin filament | 3.13 | 5.56E-04 | 1.98E-02 |  |  |  |  |
| costamere | 3.11 | 1.31E-04 | 7.60E-03 |  |  |  |  |
| dynein complex | 3.10 | 3.13E-05 | 2.97E-03 |  |  |  |  |
| viral capsid | 3.03 | 1.61E-03 | 3.93E-02 |  |  |  |  |
| apicolateral plasma membrane | 2.94 | 1.08E-03 | 3.38E-02 |  |  |  |  |
| nuclear periphery | 2.81 | 4.91E-04 | 1.88E-02 |  |  |  |  |
| viral envelope | 2.73 | 1.30E-03 | 3.52E-02 |  |  |  |  |
| desmosome | 2.65 | 5.75E-04 | 1.98E-02 |  |  |  |  |
| myosin complex | 2.46 | 3.16E-06 | 4.72E-04 |  |  |  |  |
| cell leading edge | 2.22 | 6.89E-04 | 2.30E-02 |  |  |  |  |
| adherens junction | 2.09 | 4.20E-04 | 1.67E-02 |  |  |  |  |
| kinesin complex | 2.06 | 4.00E-04 | 1.73E-02 |  |  |  |  |
| nuclear pore | 1.96 | 1.38E-04 | 7.17E-03 |  |  |  |  |
| basement membrane | 1.90 | 1.58E-04 | 7.85E-03 |  |  |  |  |
| microtubule | 1.89 | 1.55E-14 | 8.12E-12 |  |  |  |  |
| spindle pole | 1.88 | 1.14E-05 | 1.49E-03 |  |  |  |  |
| growth cone | 1.84 | 1.27E-05 | 1.47E-03 |  |  |  |  |
| centriole | 1.77 | 7.59E-05 | 4.95E-03 |  |  |  |  |
| recycling endosome | 1.69 | 4.03E-04 | 1.67E-02 |  |  |  |  |
| cytoskeleton | 1.69 | 2.80E-11 | 9.74E-09 |  |  |  |  |
| PML body | 1.67 | 1.30E-03 | 3.60E-02 |  |  |  |  |
| chromosome | 1.65 | 1.27E-03 | 3.63E-02 |  |  |  |  |
| cell-cell junction | 1.64 | 3.45E-05 | 3.00E-03 |  |  |  |  |
| dendritic spine | 1.64 | 1.94E-03 | 4.60E-02 |  |  |  |  |
| midbody | 1.62 | 5.24E-04 | 1.94E-02 |  |  |  |  |
| synapse | 1.61 | 4.95E-05 | 3.69E-03 |  |  |  |  |
| centrosome | 1.59 | 7.32E-10 | 1.91E-07 |  |  |  |  |
| axon | 1.52 | 1.09E-04 | 6.70E-03 |  |  |  |  |
| microtubule organizing center | 1.52 | 1.43E-03 | 3.75E-02 |  |  |  |  |
| actin cytoskeleton | 1.50 | 1.87E-04 | 8.45E-03 |  |  |  |  |
| cytoplasmic vesicle | 1.41 | 1.20E-03 | 3.61E-02 |  |  |  |  |
| apical plasma membrane | 1.36 | 1.45E-03 | 3.73E-02 |  |  |  |  |
| cell-cell adherens junction | 1.34 | 1.54E-03 | 3.84E-02 |  |  |  |  |
| protein complex | 1.30 | 1.20E-03 | 3.52E-02 |  |  |  |  |
| cytoplasm | 1.18 | 1.67E-15 | 1.74E-12 |  |  |  |  |
| nucleoplasm | 1.17 | 2.69E-07 | 5.62E-05 |  |  |  |  |
| membrane | 1.16 | 2.10E-05 | 2.19E-03 |  |  |  |  |
| cytosol | 1.12 | 6.14E-05 | 4.27E-03 |  |  |  |  |
| nucleus | 1.07 | 8.52E-04 | 2.74E-02 |  |  |  |  |

**Table S5:** Significant GO BP enrichments and depletions.

| Enriched |  |  |  | Depleted |  |  |  |
| --- | --- | --- | --- | --- | --- | --- | --- |
| BP | FE | p-value | Benjamini | BP | FE | p-value | Benjamini |
| tRNA export from nucleus | 2.63 | 3.69E-05 | 2.72E-02 | detection of chemical stimulus involved in sensory perception of smell | 1.20 | 6.97E-18 | 3.61E-14 |
| microtubule-based movement | 2.46 | 4.95E-10 | 1.11E-06 | detection of chemical stimulus involved in sensory perception | 1.25 | 1.37E-06 | 4.73E-03 |
| homophilic cell adhesion via plasma membrane adhesion molecules | 2.25 | 3.10E-14 | 2.09E-10 | sensory perception of smell | 1.18 | 1.24E-05 | 3.15E-02 |
| regulation of Rho protein signal transduction | 2.25 | 1.02E-07 | 1.38E-04 | G-protein coupled receptor signaling pathway | 1.15 | 8.98E-19 | 9.31E-15 |
| single organismal cell-cell adhesion | 1.98 | 2.05E-06 | 1.97E-03 |  |  |  |  |
| cytoskeleton organization | 1.76 | 1.48E-06 | 1.66E-03 |  |  |  |  |
| regulation of small GTPase mediated signal transduction | 1.72 | 3.44E-05 | 2.86E-02 |  |  |  |  |
| positive regulation of GTPase activity | 1.56 | 4.00E-12 | 1.35E-08 |  |  |  |  |
| cell adhesion | 1.52 | 8.97E-09 | 1.51E-05 |  |  |  |  |

**Table S6:** Significant GO MF enrichments and depletions.

| Enriched |  |  |  | Depleted |  |  |  |
| --- | --- | --- | --- | --- | --- | --- | --- |
| MF | FE | p-value | Benjamini | MF | FE | p-value | Benjamini |
| microfilament motor activity | 3.57 | 8.54E-07 | 1.58E-04 | odorant binding | 1.24 | 6.66E-06 | 8.48E-03 |
| structural constituent of nuclear pore | 2.97 | 1.14E-04 | 1.53E-02 | olfactory receptor activity | 1.20 | 7.19E-18 | 2.76E-14 |
| nuclear localization sequence binding | 2.71 | 7.24E-05 | 1.13E-02 | G-protein coupled receptor activity | 1.15 | 8.55E-15 | 1.64E-11 |
| microtubule motor activity | 2.68 | 2.40E-12 | 2.44E-09 |  |  |  |  |
| motor activity | 2.63 | 6.78E-10 | 3.46E-07 |  |  |  |  |
| spectrin binding | 2.57 | 4.74E-04 | 4.96E-02 |  |  |  |  |
| Rho guanyl-nucleotide exchange factor activity | 2.43 | 3.59E-09 | 1.05E-06 |  |  |  |  |
| calmodulin binding | 2.03 | 4.30E-12 | 2.92E-09 |  |  |  |  |
| ATPase activity | 1.85 | 1.25E-08 | 2.55E-06 |  |  |  |  |
| microtubule binding | 1.84 | 1.62E-09 | 5.52E-07 |  |  |  |  |
| structural constituent of cytoskeleton | 1.74 | 1.32E-04 | 1.67E-02 |  |  |  |  |
| guanyl-nucleotide exchange factor activity | 1.74 | 8.04E-05 | 1.16E-02 |  |  |  |  |
| actin binding | 1.68 | 8.75E-09 | 2.23E-06 |  |  |  |  |
| GTPase activator activity | 1.68 | 1.09E-08 | 2.47E-06 |  |  |  |  |
| protein kinase binding | 1.36 | 2.03E-04 | 2.41E-02 |  |  |  |  |
| chromatin binding | 1.35 | 3.00E-04 | 3.35E-02 |  |  |  |  |
| ATP binding | 1.34 | 1.15E-12 | 2.34E-09 |  |  |  |  |
| calcium ion binding | 1.32 | 4.90E-06 | 8.32E-04 |  |  |  |  |
| protein binding | 1.08 | 1.08E-09 | 4.42E-07 |  |  |  |  |

**Table S7:** Significant Reactome pathway enrichments and depletions.

| Enriched |  |  |  | Depleted |  |  |  |
| --- | --- | --- | --- | --- | --- | --- | --- |
| Pathway | FE | p-value | Benjamini | Pathway | FE | p-value | Benjamini |
| Cation-coupled Chloride cotransporters | 4.68 | 5.39E-04 | 2.80E-02 | Peptide chain elongation | 1.20 | 1.05E-04 | 3.80E-02 |
| Anchoring fibril formation | 4.06 | 2.07E-06 | 7.64E-04 | Viral mRNA Translation | 1.20 | 1.05E-04 | 3.80E-02 |
| Extracellular matrix organization | 3.43 | 2.00E-04 | 1.29E-02 | Formation of a pool of free 40S subunits | 1.20 | 4.62E-05 | 2.24E-02 |
| Non-integrin membrane-ECM interactions | 3.04 | 2.05E-08 | 2.27E-05 | G alpha (i) signalling events | 1.19 | 3.90E-10 | 2.88E-07 |
| Laminin interactions | 2.81 | 2.51E-05 | 3.08E-03 | Olfactory Signaling Pathway | 1.19 | 1.05E-16 | 1.64E-13 |
| Pre-NOTCH Transcription and Translation | 2.74 | 6.79E-05 | 7.49E-03 |  |  |  |  |
| NS1 Mediated Effects on Host Pathways | 2.66 | 1.27E-05 | 3.50E-03 |  |  |  |  |
| Regulation of Glucokinase by Glucokinase Regulatory Protein | 2.57 | 1.94E-04 | 1.42E-02 |  |  |  |  |
| Nuclear import of Rev protein | 2.55 | 1.26E-04 | 1.07E-02 |  |  |  |  |
| Rev-mediated nuclear export of HIV RNA | 2.48 | 2.03E-04 | 1.24E-02 |  |  |  |  |
| Vpr-mediated nuclear import of PICs | 2.41 | 4.86E-04 | 2.65E-02 |  |  |  |  |
| Nuclear Pore Complex (NPC) Disassembly | 2.41 | 3.16E-04 | 1.82E-02 |  |  |  |  |
| Assembly of collagen fibrils and other multimeric structures | 2.24 | 1.95E-04 | 1.34E-02 |  |  |  |  |
| Collagen biosynthesis and modifying enzymes | 2.17 | 1.43E-05 | 2.63E-03 |  |  |  |  |
| Loss of Nlp from mitotic centrosomes | 2.14 | 1.36E-05 | 3.00E-03 |  |  |  |  |
| SUMOylation of RNA binding proteins | 2.09 | 8.45E-04 | 3.98E-02 |  |  |  |  |
| Anchoring of the basal body to the plasma membrane | 2.05 | 1.24E-06 | 6.85E-04 |  |  |  |  |
| Recruitment of mitotic centrosome proteins and complexes | 2.04 | 2.32E-05 | 3.66E-03 |  |  |  |  |
| Regulation of PLK1 Activity at G2/M Transition | 1.97 | 2.44E-05 | 3.37E-03 |  |  |  |  |
| SUMOylation of DNA damage response and repair proteins | 1.95 | 1.26E-04 | 1.15E-02 |  |  |  |  |
| ECM proteoglycans | 1.93 | 1.84E-04 | 1.44E-02 |  |  |  |  |
| Regulation of HSF1-mediated heat shock response | 1.87 | 7.48E-04 | 3.69E-02 |  |  |  |  |
| Rho GTPase cycle | 1.74 | 8.86E-05 | 8.87E-03 |  |  |  |  |

**Table S8:**
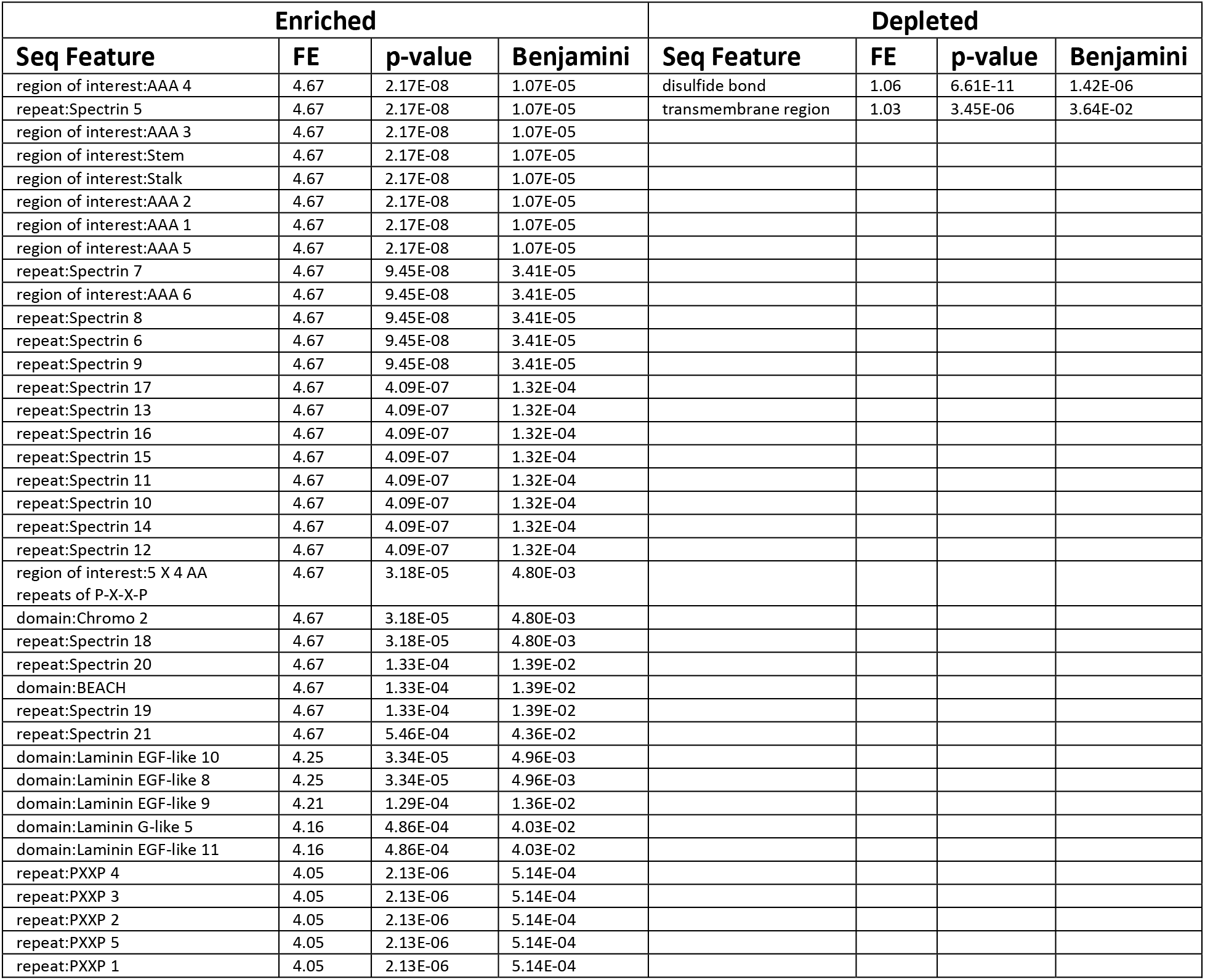
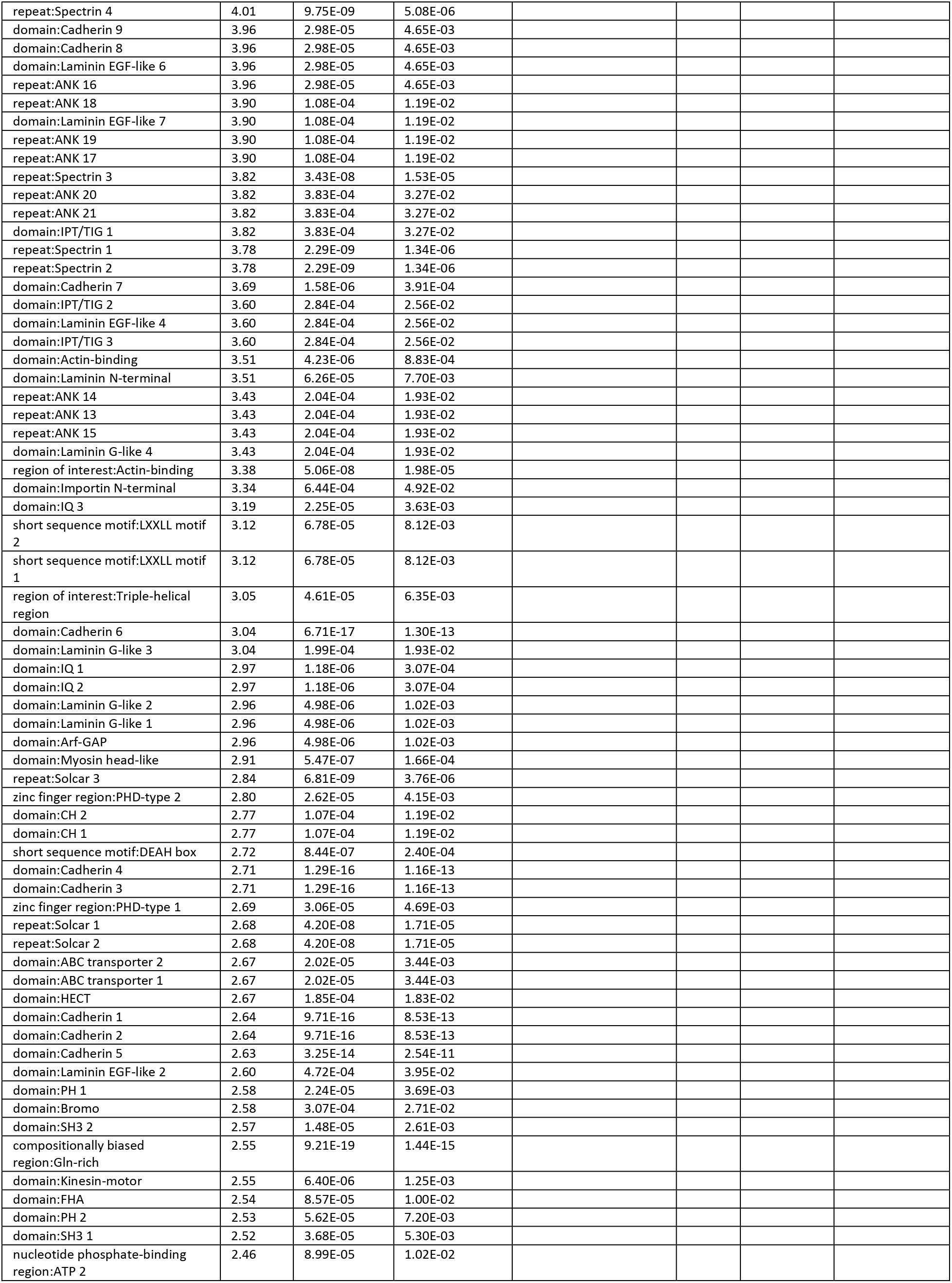
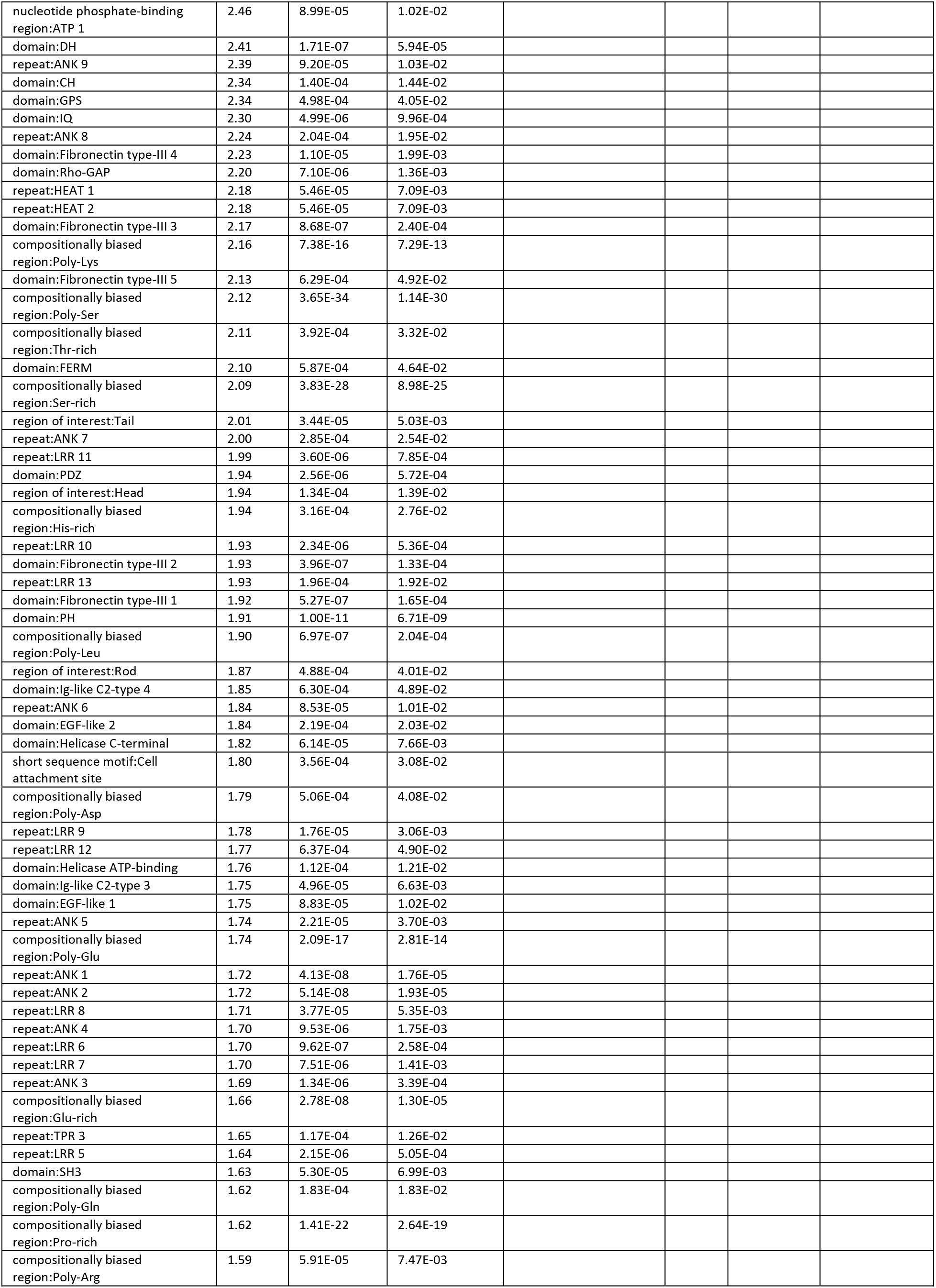
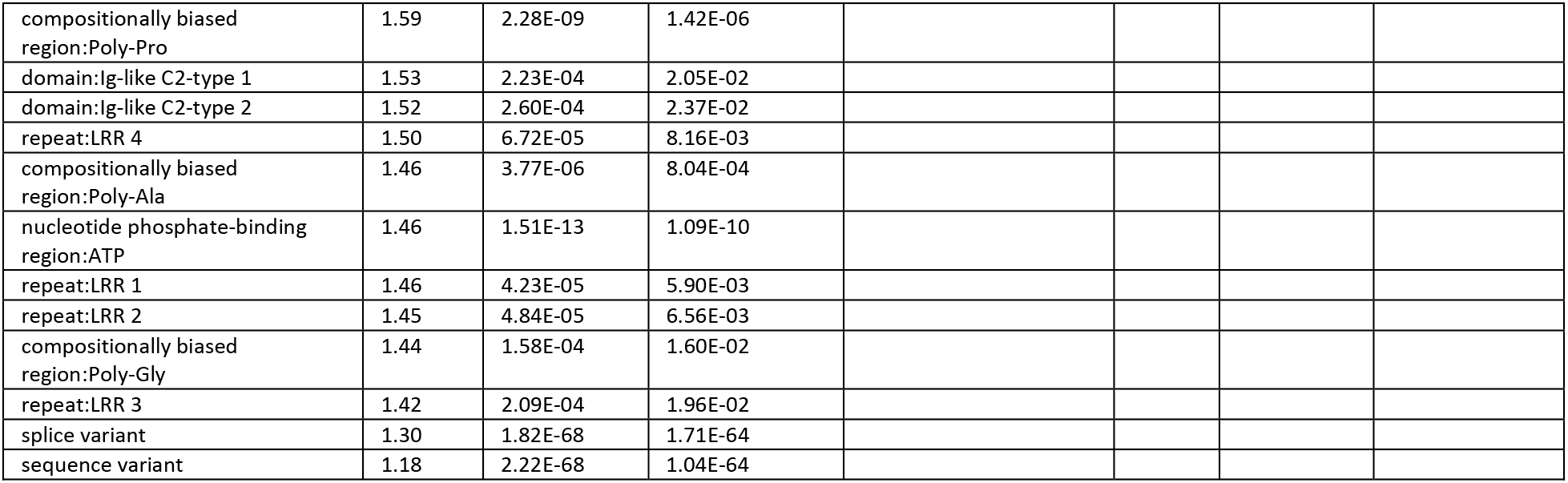
Significant sequence feature enrichments and depletions.

**Table S9:**
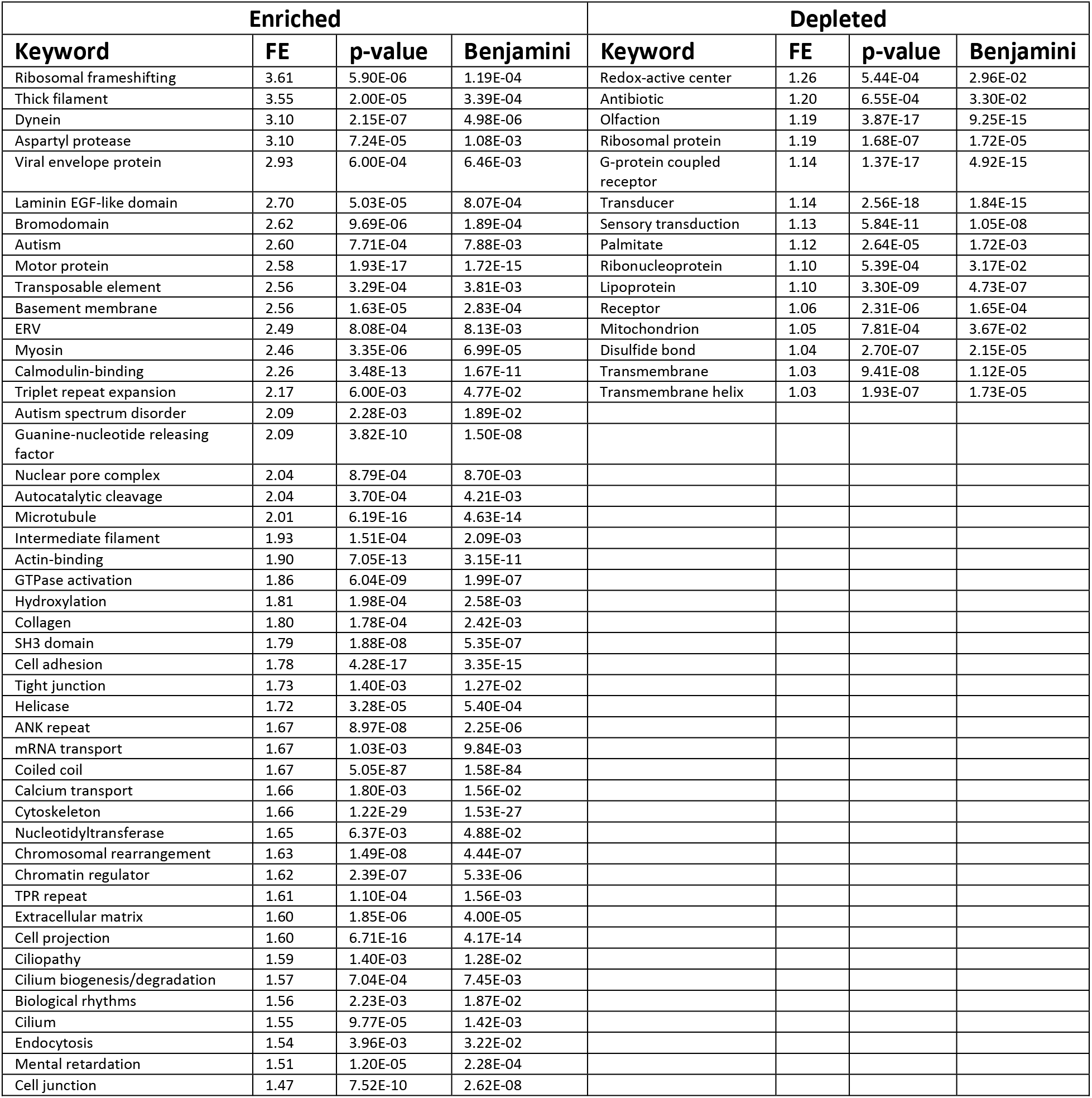
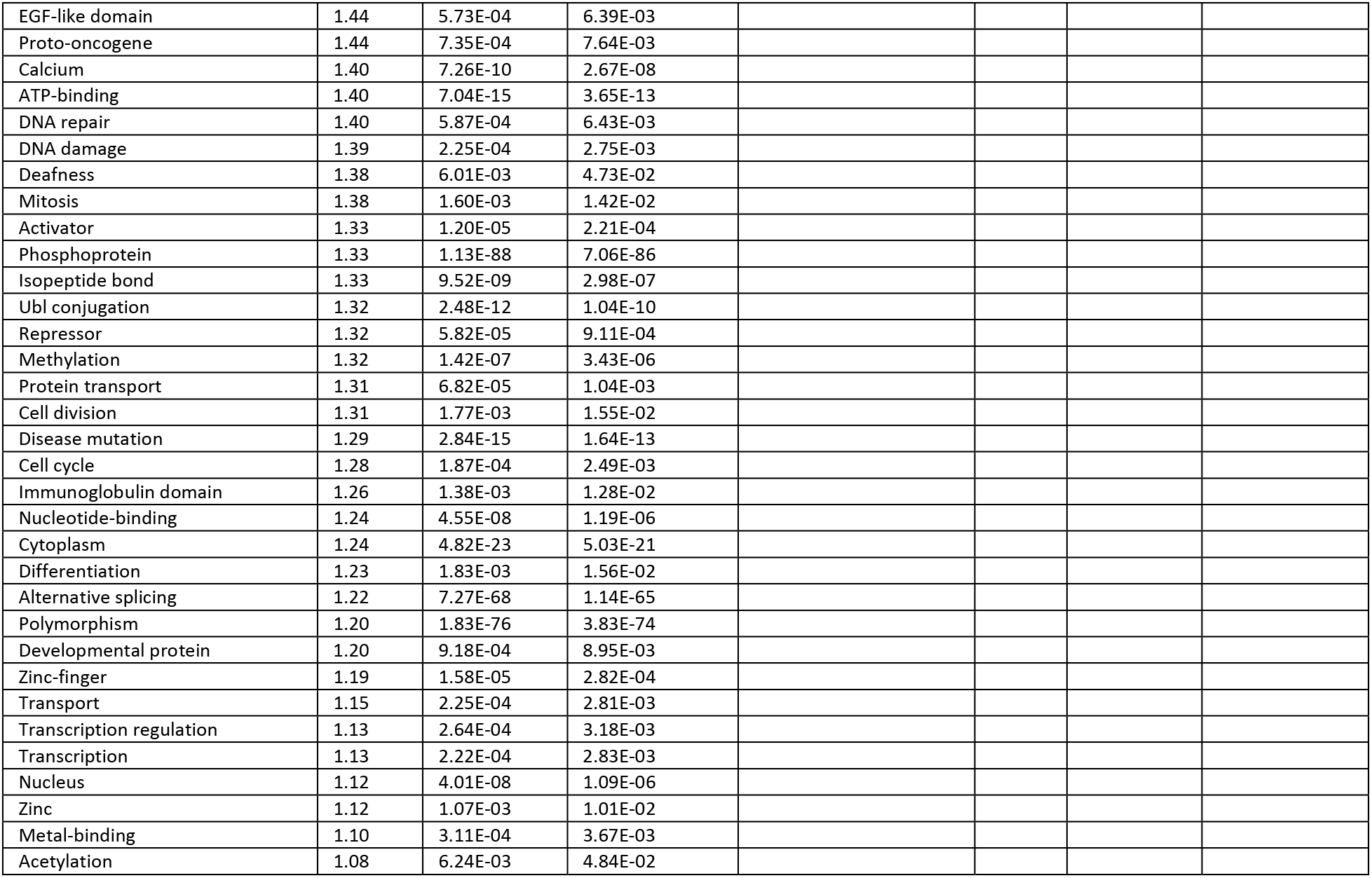
Significant keyword enrichments and depletions.

